# Nucleoid-associated proteins shape the global protein occupancy and transcriptional landscape of a clinical isolate of *Vibrio cholerae*

**DOI:** 10.1101/2023.12.30.573743

**Authors:** Yulduz Rakibova, Drew T. Dunham, Kimberley D. Seed, P. Lydia Freddolino

## Abstract

*Vibrio cholerae*, the causative agent of the diarrheal disease cholera, poses an ongoing health threat due to its wide repertoire of horizontally acquired elements (HAEs) and virulence factors. New clinical isolates of the bacterium with improved fitness abilities, often associated with HAEs, frequently emerge. The appropriate control and expression of such genetic elements is critical for the bacteria to thrive in the different environmental niches it occupies. H-NS, the histone-like nucleoid structuring protein, is the best studied xenogeneic silencer of HAEs in gamma-proteobacteria. Although H-NS and other highly abundant nucleoid-associated proteins (NAPs) have been shown to play important roles in regulating HAEs and virulence in model bacteria, we still lack a comprehensive understanding of how different NAPs modulate transcription in *V. cholerae*. By obtaining genome-wide measurements of protein occupancy and active transcription in a clinical isolate of *V. cholerae,* harboring recently discovered HAEs encoding for phage defense systems, we show that a lack of H-NS causes a robust increase in the expression of genes found in many HAEs. We further found that TsrA, a protein with partial homology to H-NS, regulates virulence genes primarily through modulation of H-NS activity. We also identified a few sites that are affected by TsrA independently of H-NS, suggesting TsrA may act with diverse regulatory mechanisms. Our results demonstrate how the combinatorial activity of NAPs is employed by a clinical isolate of an important pathogen to regulate recently discovered HAEs.

**Importance:** New strains of the bacterial pathogen *Vibrio cholerae*, bearing novel horizontally acquired elements (HAEs), frequently emerge. HAEs provide beneficial traits to the bacterium, such as antibiotic resistance and defense against invading bacteriophages. Xenogeneic silencers are proteins that help bacteria harness new HAEs and silence those HAEs until they are needed. H-NS is the best-studied xenogeneic silencer; it is one of the nucleoid-associated proteins (NAPs) in gamma-proteobacteria and is responsible for the proper regulation of HAEs within the bacterial transcriptional network. We studied the effects of H-NS and other NAPs on the HAEs of a clinical isolate of *V. cholerae*. Importantly, we found that H-NS partners with a small and poorly characterized protein, TsrA, to help domesticate new HAEs involved in bacterial survival and in causing disease. Proper understanding of the regulatory state in emerging isolates of *V. cholerae* will provide improved therapies against new isolates of the pathogen.

## Introduction

The disease cholera, caused by the bacterium *Vibrio cholerae*, occurs following the ingestion of contaminated food or water and the subsequent colonization of the small intestine by the bacterium, causing severe diarrhea induced by bacterially-produced toxins [1]. Worldwide, there are between 1.3 and 4 million annual cholera cases and an estimated 21,000 and 143,000 deaths [2,3]. *V. cholerae* is documented to rely on multiple horizontally acquired elements (HAEs) during its life cycle in both the aquatic reservoir and in human hosts [1]. HAEs provide an opportunity for bacteria to acquire beneficial traits, such as virulence factors, new metabolic properties, and antibiotic resistance via the integration of foreign DNA [4]. Both of the major *V. cholerae* virulence factors, cholera toxin and the toxin co-regulated pilus (TcpA), are encoded by genes from two distinct HAEs, the filamentous phage CTXΦ [5] and the *Vibrio* pathogenicity island-1 (VPI-1), respectively [6]. The presence of these two HAEs is a defining feature of all epidemic *V. cholerae* strains. More recently, it has been discovered that some *V. cholerae* isolates rely on an additional HAE, the phage-inducible chromosomal island-like element (PLE), to protect against infection by the highly prevalent lytic phage ICP1 [7–11]. PLE is a viral satellite, a mobile DNA element that is activated upon infection by a given phage, specifically by ICP1 for PLE, whereupon a suite of effectors deleterious to the infecting phage and a few to the host bacterium are expressed [12–14]. HAEs like PLE exemplify the trade-offs of acquiring foreign DNA for bacteria. While the contents of the PLEs provide protection against ICP1, they often do so through the expression of gene products that are toxic to the host cell (other phage-defense systems found on the HAEs have been recently investigated in *V. cholerae* [13,15–18]). Thus, erroneous transcriptional activation of HAEs may be detrimental to the host cell for different reasons, such as a waste of resources, expression of disrupted regulatory networks, or cytotoxicity, and likely would preclude stable maintenance of such HAEs unless they could be silenced under most conditions [19]. Many bacteria encode xenogeneic silencers to repress newly acquired DNA, minimizing potential harm and aiding in the domestication of new genomic elements (xenogeneic silencers and their roles in bacterial evolution are reviewed in ref. [19]).

In Gram-negative bacteria, one of the most widespread and best-studied xenogeneic silencers is H-NS [20]. H-NS is a highly abundant protein that preferentially binds and oligomerizes by forming either linear or bridged filaments on AT-rich regions (a common characteristic of HAEs [19–22]) and subsequently silences gene expression by interfering with RNA polymerase binding or promoting Rho-dependent termination via stalling or backtracking of RNA polymerase [21,23,24]. H-NS is an example of a highly abundant DNA binding protein with loose sequence specificity (a class often referred to as nucleoid-associated proteins, or NAPs, reviewed in ref. [21]) that can modulate gene expression. The model gamma-proteobacterium *Escherichia coli* has been reported to have about a dozen NAPs that change the structure of the chromatin and can act as either transcriptional activators or repressors, thus altering the entire transcriptional network in response to different conditions [21,25]. NAPs may have overlapping functions to differentially regulate diverse targets. For instance, in *E. coli*, H-NS is known to form homodimers as well as heterodimers with its paralog StpA to mediate full repression at certain loci [26,27]. H-NS can also act antagonistically with other NAPs; for example, in *E. coli*, several transcriptional units are repressed by H-NS but activated by the NAP integration host factor (IHF), a highly abundant NAP that both dramatically bends DNA and alters chromosomal supercoiling [21]. Counter-regulated H-NS/IHF targets include the adhesion-related operons *csgDEFG* and *fimB* (data from [28]). Additionally, Hha, another *E. coli* NAP, was shown to support H-NS/DNA bridged filaments despite Hha lacking a predicted DNA-binding domain [23]. While the role that NAPs play in the transcriptional regulatory landscape of *E. coli* is well established (although ongoing discoveries continue to be made on the regulatory mechanisms of NAPs in that system and other enterobacteria[23,24,27,29–32]), less is known about the mechanisms and effects of xenogeneic silencers and other NAPs in other organisms, such as *V. cholerae*.

The transcriptional regulation of many of the *V. cholerae* HAEs has been investigated, particularly those directly involved in virulence [33–35]. Previous studies of *V. cholerae* El Tor (responsible for the ongoing 7^th^ pandemic [36,37]) have shown that H-NS affects the expression of about 18% of the genome in a growth-phase-dependent manner [38]. Many H-NS targets are involved in motility, chemotaxis, biofilm development, and virulence, including the cholera toxin-encoding genes and the toxin-coregulated pilus TcpA [35,38,39]. H-NS is not the only nucleoid-associated protein implicated in regulating HAEs in *V. cholerae*. For example, IHF in *V. cholerae* has been reported to regulate the expression of virulence genes on HAEs [33] and possibly motility [40] and has been shown to be important for the conjugative transfer [41] of SXT, an integrative and conjugative element present in many *V. cholerae* strains [42,43].

Recent transcriptomic studies have demonstrated that TsrA, a protein with weak amino acid homology to H-NS, is involved in transcriptional regulation [44,45]. However, based on previous computational modeling and bioinformatics analysis, TsrA does not have a predicted DNA binding domain [44–46]. While previous studies measured the separate effects of *hns* and *tsrA* deletions on gene expression [44,45] and identified a strong overlap in the regulons of H-NS and TsrA, the combinatorial effects of H-NS and TsrA on global protein occupancy and gene expression have not yet been investigated. Additionally, the *V. cholerae* strains used in previous studies of *hns* and *tsrA* deletion did not harbor the PLE and SXT elements commonly present in clinical isolates from the current pandemic.

The transcriptional program of the phage satellite PLE has been previously interrogated by RNA sequencing during infection by ICP1 phage, revealing that PLE is transcriptionally activated upon ICP1 infection [47]. However, the factors responsible for silencing PLE in the absence of phage infection remain unidentified. The transcription of some *V. cholerae* SXT elements has been similarly interrogated, primarily through the lens of element transfer induced by antibiotics or phage infection, but not focusing on the regulatory factors responsible for silencing transcription of the SXT element [48,49]. Thus, we still lack a comprehensive understanding of the regulatory programs involving NAPs of the diverse *V. cholerae* virulence genes and HAEs shaping the bacterium’s behavior and evolution.

While the pattern of protein occupancy on the genome (from both transcription factors – TFs – and NAPs) has a profound impact on bacterial gene regulation, the large number of distinct regulatory proteins makes it impractical to separately measure the contributions of all relevant factors across biological conditions using protein-specific methods such as chromatin-immunoprecipitation followed by sequencing (ChIP-seq). The IPOD-HR (*in vivo* protein occupancy display at high resolution) method provides an alternative approach, allowing protein-agnostic measurement of the genome-wide accessibility of bacterial chromatin (**Figure 1A**) [27,50,51]. An additional parallel step during IPOD-HR involves ChIP-seq of RNA polymerase-bound DNA regions, which provides information on RNA polymerase (RNAP) occupancy, a proxy for active transcription [51], and allows for the distinction of transcribed vs. silenced genomic contexts. The IPOD-HR workflow has been previously applied to *E. coli* and *Bacillus subtilis* to investigate their transcriptional programs and the effects that certain regulatory factors (especially NAPs) have on genome-wide protein occupancy, chromatin accessibility, and transcription [27].

**Figure 1.**
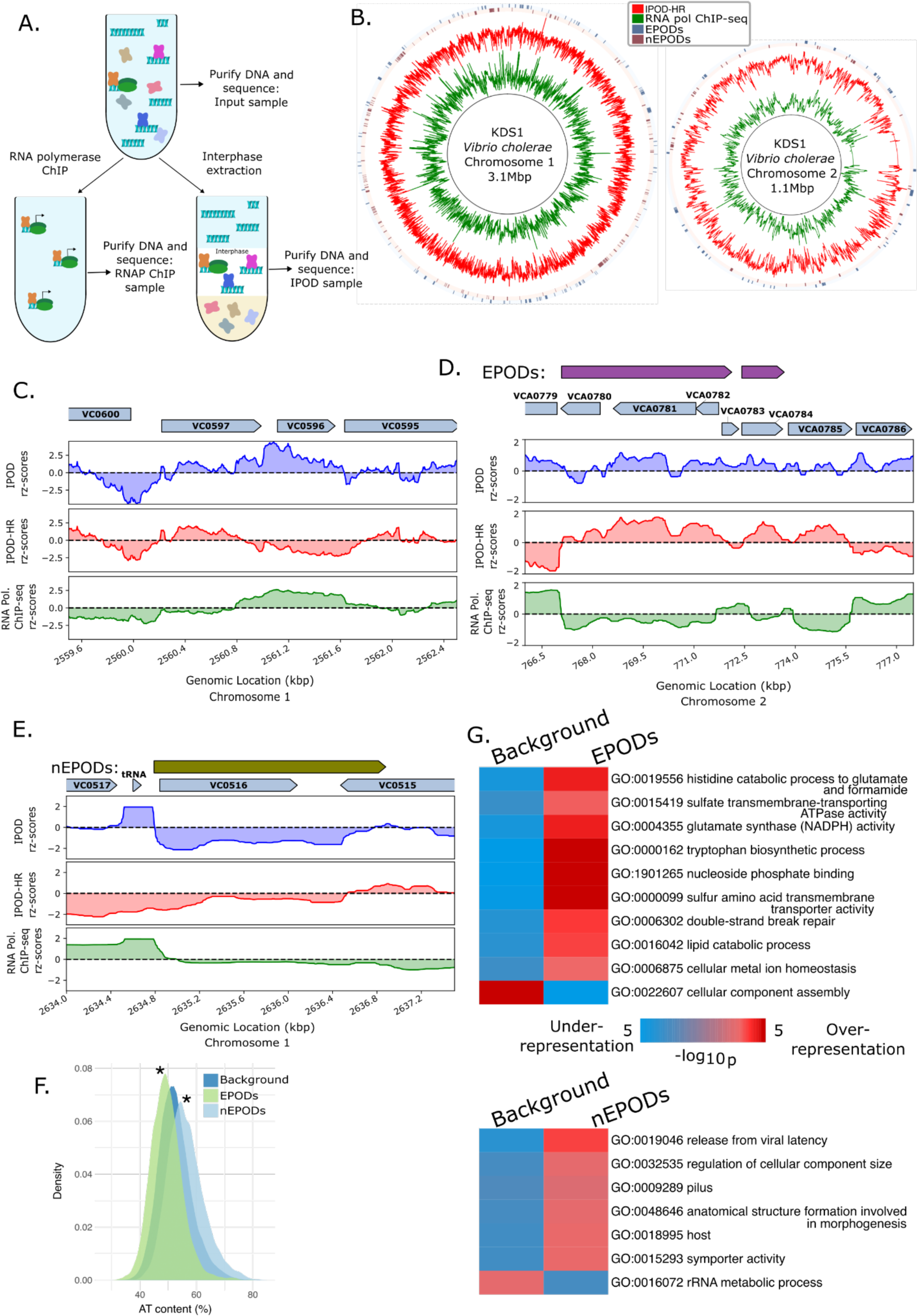
(previous page): Overview of the protein occupancy landscape in *Vibrio cholerae*. A) Experimental diagram of the IPOD-HR workflow: crosslinked protein-DNA complexes are divided into three samples: “Input” represents the baseline total fragmented DNA, which is then subjected to extraction of the protein-DNA complexes enriched in the interphase layer (IPOD) and, in parallel, to chromatin immunoprecipitation (ChIP) with an anti-RNA polymerase antibody [64]. DNA from the three samples is purified and sequenced. DNA is depicted in light blue, RNA polymerase (RNAP) is depicted in green with two lobes, the rest of the proteins in the samples are shown as x-shaped cartoons. B) Circos [65] plots of the *V. cholerae* KDS1 genome that include IPOD-HR traces of rz-scores, RNA polymerase ChIP-seq rz-scores, and locations of EPODs and nEPODs (data tracks are rolling medians over 512 bp windows). C) An example of likely local regulator binding in wild type *V. cholerae*; shown are typical binding patterns of DNA-binding regulatory proteins (through the midst of VC0597) and of highly transcribed regions (VC0596). IPOD rz-scores are based on the log2 ratios of extracted/input DNA. IPOD-HR rz-scores are RNAP ChIP subtracted IPOD rz-scores (see Materials and Methods). RNA polymerase ChIP-seq rz-scores are based on the log2 ratio of extracted RNAP ChIP vs input DNA. The VC numbers for genes from the C6706 strain of *V. cholerae* have been mapped to the KDS1 strain used in this study. D) Example of a transcriptionally silent extended protein occupancy domain (EPOD) in wild type *V.cholerae,* showing high protein occupancy and low RNA polymerase ChIP-seq signal across two divergent operons (spanning VCA0780-VCA0785). For this panel and the rest of the manuscript, the plotted rz-scores are median rz-scores over 512 bp windows unless otherwise noted. E) Example of a negative EPOD (nEPOD) in *V. cholerae,* showing both negative IPOD enrichment scores and depleted RNA polymerase ChIP-seq signal across VC0516 and partially on VC0515. F) Distribution of AT content of EPOD regions in the wild type *V. cholerae* KDS1, where negative EPODs (nEPODs) are higher in AT content than the background, whereas positive EPODs have higher GC content. “Background” is composed of all genomic regions that do not fall into the other indicated categories. Significance calling was performed with 1,000-sample permutation tests (described in detail in Materials and Methods) of differences in medians of AT content between the background and the EPODs or nEPODs; in both cases, a significant difference was observed (p-value=0.001). G) GO term enrichment analysis of EPODs and nEPODs. In the iPAGE plots in this manuscript, the color of the heatmap indicates the degree of enrichment or depletion of a given GO term (see color scale) among the genes in each GO term (row), in the EPODs or nEPODs under consideration vs. the rest of the genome.

In order to comprehensively determine the roles of key NAPs in regulating the HAEs of *V. cholerae*, we employed IPOD-HR together with RNA sequencing on the *V. cholerae* clinical isolate KDS1, as well as *Δhns*, *ΔtsrA*, *ΔihfA* (IhfA subunit of IHF), and *ΔtsrAΔhns* deletion mutants derived from the same parental strain. Consistent with previous findings, we observed that the absence of H-NS altered protein occupancy and gene expression in many genomic regions important for host colonization and virulence, including multiple HAEs. We also show that the deletion of *hns* leads to the de-repression of some portions of the phage satellite PLE, suggesting that phage infection perturbs H-NS occupancy, thus contributing to the activation of the PLE program. Analysis of the *ΔtsrAΔhns* double knockout strain revealed that H-NS drives the main repression of targets and that TsrA generally acts in an H-NS-dependent manner; however, we also identified some sites where TsrA exerts H-NS independent regulatory control. Given the importance of the SOS response in inducing mobile genetic elements, such as prophages and antibiotic resistance genes [52–54], we also explored the effect of DNA damage on protein occupancy profiles and gene expression by performing IPOD-HR on *V. cholerae* treated with the DNA damaging agent mitomycin C (MMC). We observed a bimodal response in HAEs following MMC treatment: MMC induces increased transcription of some HAEs but increased silencing of others, likely due to substantial rearrangements of the chromatin state at those loci. Our findings both reaffirm prior knowledge regarding the importance of NAPs (especially H-NS) for regulating HAEs in a clinical isolate of *V. cholerae* and reveal the combinatorial regulation between NAPs and other local regulators.

## Results

### IPOD-HR identifies large repressive protein occupancy regions in addition to local protein binding in *Vibrio cholerae*

In order to identify locations of potentially repressive NAP binding genome-wide in *V. cholerae*, we applied IPOD-HR (*in vivo* protein occupancy display at high resolution) to *V. cholerae* KDS1 cells grown to exponential phase in lysogeny broth (LB). KDS1 is a clinical isolate representative of the *V. cholerae* O1 serogroup El Tor biotype, isolated in Bangladesh in 2011 [55], and will be referred to simply as *V. cholerae* hereafter. As detailed in [50,51], IPOD-HR provides a snapshot of total protein occupancy along the genome, along with paired RNA polymerase occupancy data (**Figure 1A**). Throughout the text below, we refer to the RNAP ChIP-seq subtracted protein signal as IPOD-HR, and the total protein occupancy before RNAP subtraction as IPOD.

Application of IPOD-HR to *V. cholerae* revealed trends that are consistent with those identified in other model organisms [27,50,51]. The *V. cholerae* genome shows a diverse protein occupancy pattern, including local regions of protein occupancy consistent with transcription factor binding sites and large transcriptionally silent regions consistent with extended regions of NAP occupancy referred to as extended protein occupancy domains (EPODs), which have been observed in other organisms [27,50,51] (**Figure 1B**). An example of the information provided by IPOD-HR at a single locus can be seen in **Figure 1C**, where VC0596 (encoding *dksA*, a transcription factor [56]) appears highly expressed based on high RNA polymerase (RNAP) occupancy throughout the open reading frame, whereas the adjacent gene VC0597 (*sfsA*, annotated as “sugar fermentation stimulation protein homolog” in Uniprot) lacks RNAP occupancy and appears to be bound by a local regulator that represses VC0597 and/or activates transcription of VC0596. An example of a large EPOD is seen in **Figure 1D**, where a region of high protein occupancy extends about 8 kilobases over the region of VCA0780-VCA0785; RNA polymerase binding is generally reduced over the same region. Of the genes encompassed by this EPOD, VCA0785 encodes for CdgC, which acts as a c-di-GMP synthase/phosphodiesterase and is known to regulate biofilm formation and motility, as well as virulence factor expression and rugose colony morphology [57,58]. The rest of the ORFs in this EPOD have not been characterized and have only automatically inferred annotations [59,60]. As was shown in the application of IPOD-HR in *E. coli* [27,50,51], EPODs tend to be transcriptionally silent, as is seen for the VCA0780-VCA0785 EPOD under the growth condition considered here.

Previous application of IPOD-HR to *B. subtilis* revealed that some EPODs, such as those representing occupancy by the major NAP Rok, show negative enrichment scores following IPOD interphase extraction, in contrast to what was observed for the major NAPs in *E. coli* [27]. The most likely explanation is Rok-DNA complexes partitioning away from the interphase layer due to the properties of the Rok protein, and thus being depleted during the phenol-chloroform extraction rather than enriched, resulting in a negative ‘enrichment’ score. In considering protein occupancy across the *V. cholerae* chromosome, we observed regions of strong, sustained negative IPOD occupancy (before and after RNA polymerase ChIP-seq subtraction) similar to those observed in Rok-occupied regions in *B. subtilis*; an example is shown in **Figure 1E**. Importantly, these large negative occupancy regions appear to be transcriptionally silent, which is consistent with repressive NAP occupancy, and indicates that the observed negative signal is not due to the RNAP ChIP-seq subtraction used in the IPOD-HR method, or due to unbound DNA, which has been accounted for through comparison with the input sample. We refer to regions of depleted IPOD signal and low expression as negative EPODs (nEPODS) (**Figure 1B**). In the case of the example region shown in **Figure 1E**, we see a 2 kilobase nEPOD encompassing VC0516 and partially VC0515 on the large chromosome of *V. cholerae*. The region is a part of the HAE *Vibrio* seventh pandemic island-II (VSP-II) [61], where the transcriptionally silent VC0516 corresponds to the phage-like integrase of the VSP-II [62]. We hypothesize that, as in the *B. subtilis* Rok case noted above, the negative occupancy signal represents a protein or a combination of proteins that produce this behavior due to their properties in the IPOD-HR protocol. As detailed in the Supplementary Text, given the high overlap between known H-NS bound regions and nEPODs, we hypothesize that many of the nEPODs in *V. cholerae* may represent H-NS binding.

In *E. coli*, EPODs have been observed to form mainly on AT-rich DNA, and to regulate many horizontally acquired genes, prophages, and mobile genetic elements [27,50,51], consistent with previously established regulatory roles of xenogeneic silencers [19–22]. To characterize the properties of EPODs and nEPODs in *V. cholerae*, we examined the AT richness of EPODs, as well as performed gene set enrichment analysis to identify gene ontology (GO) terms occurring frequently in EPODs. In contrast to *E. coli* [27,50,51], EPODs in *V. cholerae* are significantly higher in GC content compared with a background corresponding to the rest of the genome, whereas nEPODs are significantly higher in AT content (**Figure 1F**). These results suggest that EPODs and nEPODs in *V. cholerae* likely correspond to occupancy from different DNA binding proteins (as previously observed in *B. subtilis*). Both EPODs and nEPODs, however, are associated with transcriptional silencing represented by a lack of RNAP occupancy in those regions. In order to identify the pathways primarily affected by EPODs in *V. cholerae*, we utilized the iPAGE method for gene set enrichment analysis [63], which identifies significant correspondences between gene ontology (GO) terms and the EPOD status (via mutual information) (**Figure 1G**). Primarily metabolism-oriented GO terms, such as “histidine catabolic process to glutamate and formamide” and “tryptophan biosynthetic process”, are over-represented in positive EPODs, and terms such as “release from viral latency” and “pilus” are enriched in negative EPODs (nEPODs), suggestive of HAE-related processes appearing in nEPODs. Collectively, the enriched GO term categories in *V. cholerae* EPODs and nEPODs match with the categories enriched in EPODs in *E. coli,* covering a spectrum of specialized metabolic functions and horizontally acquired elements. The distinction in categories between EPODs and nEPODs likely reflects the division of regulatory labor between different NAPs in *V. cholerae,* as was observed in *B. subtilis*.

### Nucleoid-associated protein IHF in *Vibrio cholerae* is involved in global iron regulation

Due to the many transcriptional regulators in a bacterial cell, the IPOD-HR method allows us to identify regulatory and protein occupancy effects along the genome upon the deletion of a NAP from both direct and indirect binding of effects of the NAP. The integration host factor (IHF) is one of the NAPs that acts as a dual regulator, broadly affecting gene expression [21]. Thus, we profiled the transcriptional network of *V. cholerae* lacking the IhfA subunit, which dimerizes with the IhfB subunit to make functional IHF. With IPOD-HR we identified **(Table S1)** distinct loci at which protein occupancy is lost in the absence of IHF compared to wild type (**Figure 2A**). One such example occurs in the region upstream of VC0143, a hypothetical protein predicted to be essential due to the lack of transposon insertions in whole-genome transposon library screens [66]. We observe a decrease in protein occupancy coupled with an increase in RNA polymerase occupancy in the VC0143 promoter region upon *ihfA* deletion (**Figure 2A**). The intergenic region between VC0142 and VC0143 has been reported to encode a small RNA [67,68]. It was also shown that this region contains binding sites for the ferric uptake regulator (Fur) based on ChIP-seq data [67] and for the virulence regulator ToxT based on *in vitro* DNA pull-down of purified ToxT [69]. Thus, previous studies suggest that the expression of the VC0142/VC0143 sRNA may be regulated by multiple factors, including: Fur [67], which is known to respond to intracellular iron levels; ToxT [69], the master virulence regulator; and the nucleoid-associated protein IHF either directly or indirectly (based on our data). In order to trace the path from *ihfA* deletion to VC0142/VC0143 sRNA expression, we considered how the Fur regulon is, as well as iron homeostasis-related genes, are affected by the absence of IHF. RNA-sequencing (RNA-seq) showed that many genes involved in iron ion homeostasis and/or known to be in the Fur regulon are upregulated in the absence of IHF (**Figure 2B**). Our RNA-seq data also corroborated the findings from IPOD-HR that in the absence of IHF, the sRNA in the VC0142/VC0143 intergenic region as well as the hypothetical protein VC0143 are both significantly upregulated (log2 Fold Change (FC) and q-value of 3.94/1.01e-09 and 5.22/3.21e-14 for the small RNA and VC0143, respectively). iPAGE GO term enrichment analysis shows that the “iron ion transport” and “siderophore-biosynthesis process” are some of the enriched GO terms for highly expressed genes, consistent with the above findings that many iron ion homeostasis genes are upregulated in the absence of the alpha subunit of IHF (**Figure 2C, Supplementary Figure 1A**).

**Figure 2.**
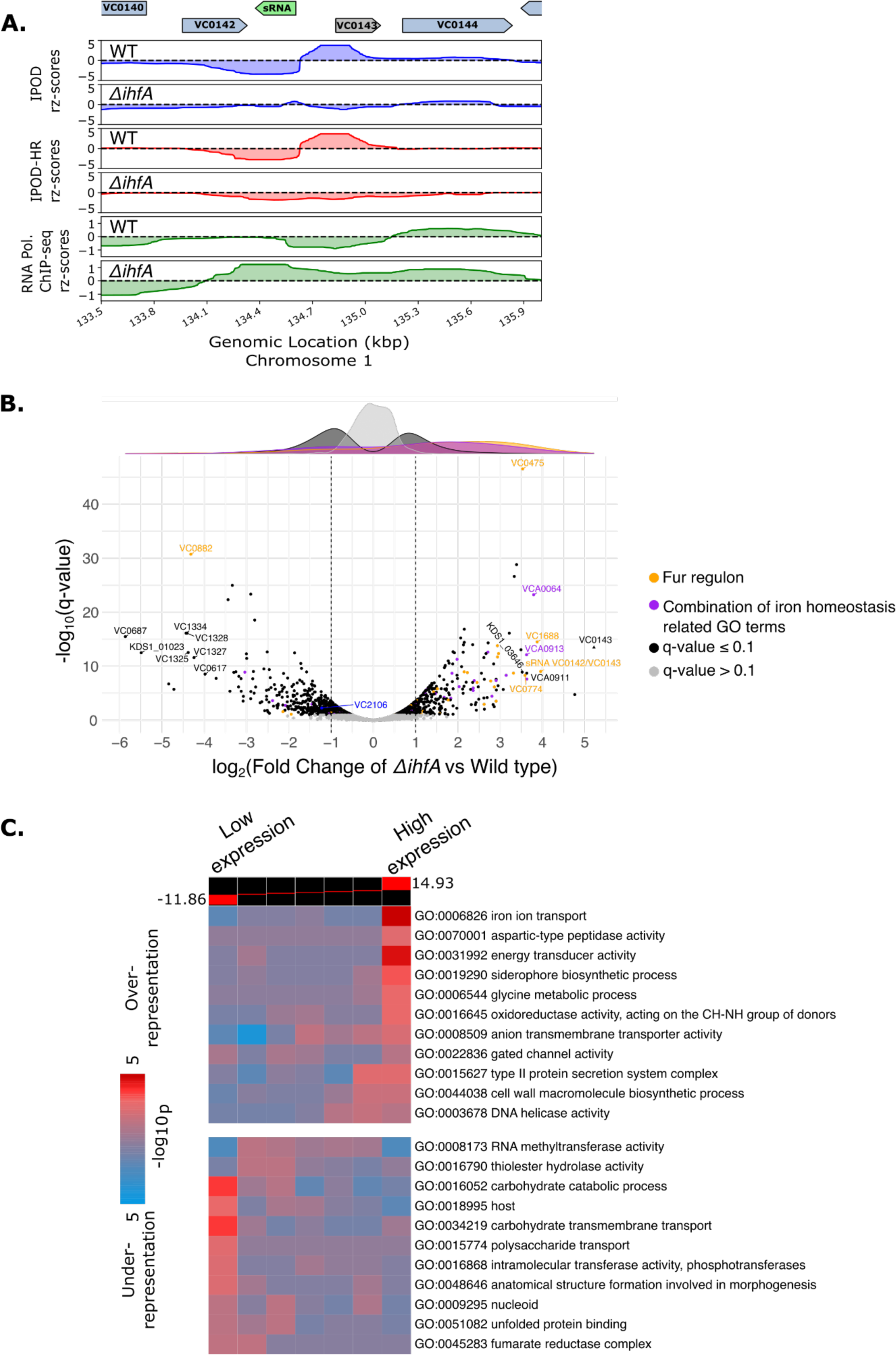
(previous page): Protein occupancy and RNA-sequencing in *V. cholerae* lacking IHF. A) Protein occupancy (total and RNA polymerase) in the promoter region upstream of VC0143 and the VC0142/VC0143 intergenic sRNA for WT and *ΔihfA* cells during exponential growth in LB media. The sRNA is shown in green and VC0143 is shown in gray because the annotation pipeline utilized only annotated genes coding for more than 90 amino acids (see Materials and Methods), and thus manual annotation was required. B) Volcano plot of differentially expressed genes in the strains lacking IHF (*ΔihfA*) relative to wild type cells. Dark colored points above the gray points represent genes that passed the significance threshold of q-value less than or equal to 0.1. The *fur* (VC2106) gene is denoted in blue, Fur regulon genes obtained from the ChIP-seq results found in ref. [67] (with q-value less than or equal to 0.1) are represented in orange, and a combination of GO terms that had iron in their names are represented in purple. Marginal density plots follow the same color coding as the points in the volcano plot. VC numbers (mapped from C6706 strain to KDS1 *V. cholerae*) or KDS1 numbers (that did not map with VC numbers) are only shown for highly (log2FC greater or equal to 3.5 or −3.5) differentially expressed genes, or the VC0142/VC0143 sRNA and the *fur* gene, to avoid crowding. VC0143 is manually annotated and shown as a triangle shape. C) GO term enrichment analysis of differentially expressed genes in the *ΔihfA* strain (relative to WT) from RNA-sequencing via iPAGE. Wald statistics from the *ΔihfA* RNA-seq are utilized as the input for iPAGE, detailed in Materials and Methods. The colors on the heatmap represent the GO terms that are highly expressed in *ΔihfA* (right) or repressed (left).

Given the upregulation of iron homeostasis-associated genes in the absence of IhfA, we assessed Fur (VC2106) expression and observed a log2FC of −1.22 (q-value=0.005) in the *ΔihfA* strain compared to wild type, suggesting that Fur expression is decreased upon loss of functional IHF, providing a ready mechanism connecting loss of IHF to the widespread changes in the regulation of iron uptake that we observed.

Interestingly, Fur is also downregulated in the knockout strains mentioned below (*ΔtsrAΔhns* (L2FC=-2.71/q-value=1.08e-21)*, ΔtsrA* (L2FC=-1.54/q-value=6.83e-07) and *Δhns* (L2FC=-1.69/q-value=2.60e-06)), but we did not observe a large global effect on the expression of iron homeostasis genes as observed in the *ΔihfA*, suggesting that IHF may be playing a distinct role in regulating the genes involved in iron-homeostasis rather than acting only through Fur.

### The absence of H-NS results in increased RNA polymerase occupancy across horizontally acquired elements in *V. cholerae*

In order to investigate the global regulatory effects of the xenogeneic silencer H-NS, as well as the putative H-NS-associated regulatory factor TsrA [46] (the regulon of which overlaps with that of H-NS [44,45]), we applied a combination of IPOD, RNA polymerase ChIP-seq, and RNA-seq to unravel changes in protein occupancy and gene expression in the absence of H-NS, TsrA or both. We compared the results to the effects of deleting *ihfA* (described above), and to a deletion of the enterobactin receptor gene *vctA* [70], which is not expected to have substantial effects under the growth conditions considered here, but serves as a control for any effects of constructing the deletions themselves.

To quantify the effects of the NAP deletions indicated here on regions of likely NAP-mediated silencing, we calculated the changes in occupancy (relative to WT) for different deletion strains at the locations of EPODs and nEPODs identified in our wild type strain **(Supplementary figure 2**). We observe that, while many EPODs are unaffected, a substantial subset show very strong loss of occupancy (based on negative tails in the IPOD-HR violin plots) in the *Δhns, ΔtsrAΔhns, ΔtsrA,* and *Δihf* strains. Similarly, while RNAP occupancy is in the average case unchanged, a subset of EPODs and nEPODs show a sharp increase in RNAP occupancy in the *Δhns* and *ΔtsrAΔhns* strains, indicating that deletion of *hns*, but not of the other regulators considered here, is sufficient to trigger expression changes in those regions. This observation indicates that the selected NAPs result in changes in protein occupancy in EPODs/nEPODs, but only the lack of H-NS results in significant de-repression of those regions. Deletion of the *vctA* control gene shows minimal shifts in the centers of the occupancy distributions relative to WT, but in the opposite direction as any NAP deletion, and without any substantive tail that would indicate strongly affected specific loci.

Because some NAPs like H-NS are involved in repression of HAEs, we investigated protein occupancy in some of the known *V. cholerae* HAEs: *ctx* and the RS1 phage satellite, the *Vibrio* pathogenicity islands (VPI-1 and VPI-2), the *Vibrio* seventh pandemic islands (VSP-I and VSP-II), the integrative and conjugative element SXT-*Vch*Ind6, the viral satellite PLE, and the superintegron (**Figure 3A**). We find that the absence of H-NS (in both the *Δhns* and *ΔtsrAΔhns* strains) results in strong de-repression in VPI-1 (**Figure 3C**), PLE (**Figure 3E**), and some de-repression in CTX (**Figure 3D**), VPI-2, VSP-I, VSP-II based on RNA polymerase ChIP-seq (**Figure 3B, Supplementary Figure 3,4**). Deletion of *tsrA* only largely affected the transcription in VPI-1 out of the considered HAEs **(Supplementary Figure 4C**). Consistent with previous H-NS ChIP-seq results [71] (**Figure 3C, gray track)** showing that H-NS covers almost the entirety of VPI-1, we observe that the absence of H-NS results in de-repression of VPI-1 as evidenced by the appearance of a large stretch of RNA-polymerase occupancy in the *Δhns* and *ΔtsrAΔhns* strains (**Figure 3C; Supplementary Figure 5A**). As with VPI-1, deletion of *hns* results in de-repression of *ctxA* and *ctxB,* but no significant de-repression is observed in the other deletion strains tested here (**Figure 3D, Supplementary Figure 4C, 5C)**.

**Figure 3.**
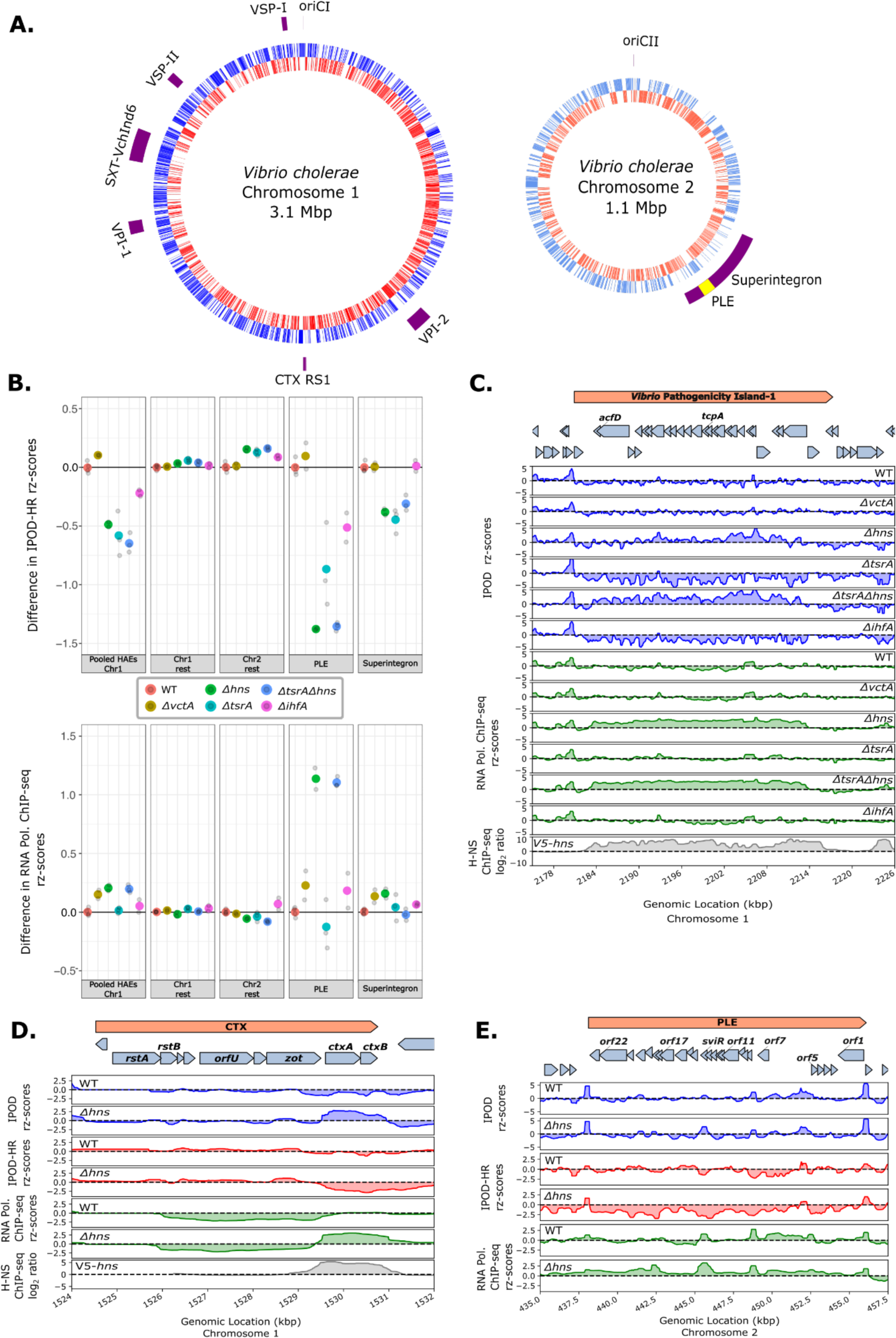
(previous page): Effect of NAP deletions on the horizontally acquired elements of *V. cholerae*. A) Locations of known HAEs in the KDS1 strain of *V. cholerae* (outer rings). The blue and orange lines in both chromosomes indicate plus and minus strand genes, respectively. The two chromosomes are not depicted to scale. B) Changes (relative to WT) in robust rz-scores of IPOD-HR and RNA polymerase occupancy across the indicated genomic regions. Here and in subsequent panels summarizing IPOD-HR occupancies, a 50 bp rolling median was used as the fundamental unit of data for each genomic location, and the plotted values reflect the pseudomedian of those values across the indicated genomic features; for RNA polymerase ChIP-seq, we followed a similar procedure, except that we used gene-level means of the ChIP occupancies as individual units of data for the pseudomedian calculations. Individual biological replicates are shown in gray points and the larger colored points are the mean of replicates for each genotype. C) Total protein occupancy signals and RNA polymerase occupancy in VPI-1 for all of the genotypes studied. The gray track shows previously obtained V5-HNS ChIP-seq from C6706 [71] remapped and requantified on our reference genome (detailed in Materials and Methods).The tracks of total protein occupancy with RNAP ChIP subtraction (IPOD-HR) are found in Supplementary Figure 5A. D) As in panel C, for the CTX prophage region; the additional red tracks indicate IPOD-HR. Only wild type and *Δhns* are depicted in Figure 3D, the rest of the genotypes are found in Supplementary Figure 5C. E) Total protein occupancy signals and RNA polymerase occupancy in the PLE satellite in wild type and *Δhns V. cholerae*. The rest of the genotypes are found in Supplementary Figure 5D.

Although H-NS regulation of VPI-1 and CTX have been studied in other *V. cholerae* strains [35,38], the regulatory factors that modulate some of the recently identified HAEs, like the phage satellite PLE, in different clinical isolates have not been investigated. In the absence of H-NS, the later ORFs in PLE become de-repressed as evidenced by the observed substantial increase in RNA polymerase occupancy coinciding with the SviR promoter (**Figure 3E, Supplementary Figure 5D**). This is consistent with SviR expression being below the limit of detection in the absence of ICP1 infection [72] and the absence of H-NS allowing for transcription from the SviR promoter and RNA polymerase occupancy extending into downstream ORFs.

Additionally, no dramatic changes are observed across PLE in RNA polymerase occupancy in the absence of *tsrA, ihfA,* or the control *vctA*, suggesting that H-NS is the primary repressor of PLE gene expression. However, the RNA polymerase occupancy in the absence of H-NS is inconsistent with the highly programmed gene expression profile of PLE responding to ICP1 infection, which is known to activate the PLE transcriptional program [47]. The incomplete activation of PLE gene expression is consistent with previous observations that other regulatory factors (presumably ICP1-derived) are needed to fully activate the PLE. Our IPOD-HR results show that among the tested NAPs, H-NS results in the highest level of repression of the *V. cholerae* HAEs, and that H-NS repression plays a considerable role in silencing aberrant PLE gene expression in the absence of ICP1 infection.

Although there is a high correlation between the RNA polymerase ChIP-seq and RNA-seq results of our *V. cholerae* strain **(Supplementary Figure 8A**), we further assessed the effect sizes of fold changes in transcript levels of individual genes in the PLE with RNA-seq. We observed that *orf15*, coding for a nicking endonuclease, NixI, that cleaves and inhibits the ICP1 phage genome replication [13], is the most highly differentially expressed gene in cells lacking *hns* (with L2FC= 5.29/q-value=6.52e-19) out of the all annotated genes in the PLE (**Table 1**). Additionally, *orf17/tcaP* is also the most significantly differentially (with L2FC=3.11/q-value=5.09e-35) expressed gene that was recently demonstrated to encode a scaffolding protein of the phage ICP1 coat allowing better transmission of the PLE HAE [14]. Consistent with our findings that the loss of *hns* masks the effect of the loss of *tsrA*, H-NS is silencing the majority of the PLE (**Table 1**) and no further effect is seen when both H-NS and TsrA are absent. Although lack of H-NS results in de-repression of the majority of genes in the PLE, we did not observe the excision or replication of the phage satellite (based on the read counts of the PLE in the input samples of strains lacking *hns*, and a lack of read boundaries at the PLE ends that would indicate excision). This is consistent with ICP1-encoded PexA being necessary for excision of PLE [11]. Taken together, our IPOD-HR and RNA-seq results show that out of the tested NAPs, H-NS repression plays a considerable role in silencing aberrant PLE gene expression in uninfected cells.

**Table 1:** RNA-sequencing Log2 (Fold Change (FC)) of deletion strains (relative to wild type) in genes that have been annotated in the PLE of *V. cholerae*. We consider a q-value of ≤ 0.1 to be significant. The genes have been sorted by increasing q-values in the *Δhns* vs wild type. *gene was manually annotated.

| PLE genes | Log2FC in $\Delta hns$ vs wild type | q-value in $\Delta hns$ vs wild type | Log2FC in $\Delta tsrA$ vs wild type | q-value in $\Delta tsrA$ vs wild type |
| --- | --- | --- | --- | --- |
| ORF17 (TcaP) | 3.113 | 5.09E-35 | 0.686 | 6.97E-03 |
| ORF15 (NixI) | 5.289 | 6.52E-19 | 2.287 | 6.28E-05 |
| ORF13 | 2.049 | 1.55E-15 | 1.796 | 3.36E-15 |
| ORF2 (CapR) | 2.424 | 4.74E-15 | 2.320 | 1.62E-17 |
| ORF21 | 1.885 | 2.77E-11 | 0.366 | 2.09E-01 |
| ORF16 | 3.541 | 3.86E-09 | 0.293 | 6.69E-01 |
| ORF15.1 | 3.806 | 1.22E-07 | -0.149 | 8.69E-01 |
| ORF14 | 1.836 | 5.19E-07 | 1.849 | 4.15E-09 |
| ORF22 | 1.598 | 8.79E-06 | -0.152 | 7.02E-01 |
| ORF3 | 2.229 | 2.53E-05 | 1.360 | 3.78E-03 |
| ORF23 | 1.347 | 5.73E-05 | -0.225 | 5.18E-01 |
| ORF12 | 1.658 | 5.13E-04 | 1.111 | 7.28E-03 |
| ORF9 | 1.104 | 5.56E-04 | 1.068 | 7.55E-05 |
| ORF5 | 2.436 | 1.36E-03 | 2.599 | 4.07E-05 |
| ORF4 | 1.946 | 3.86E-03 | 0.836 | 1.64E-01 |
| ORF8 | 0.764 | 5.35E-03 | 1.020 | 3.66E-06 |
| SviR (sRNA) | 1.414 | 2.79E-02 | 1.465 | 4.34E-03 |
| ORF18.1 | -1.095 | 3.85E-02 | -1.547 | 1.59E-04 |
| ORF20.1 (LidI)* | 1.039 | 4.45E-02 | -0.685 | 1.09E-01 |
| ORF7 | -1.015 | 5.88E-02 | -1.096 | 8.87E-03 |
| ORF18 | -0.767 | 1.74E-01 | -1.162 | 5.46E-03 |
| ORF12.1 | 1.321 | 2.50E-01 | 0.521 | 5.98E-01 |
| ORF11 | 0.615 | 2.68E-01 | -0.176 | 7.19E-01 |
| ORF19 | -0.181 | 5.93E-01 | -0.659 | 1.85E-03 |
| ORF20 | 0.306 | 6.13E-01 | -1.234 | 9.27E-04 |
| ORF1 | -0.240 | 6.34E-01 | -0.268 | 4.53E-01 |
| ORF10 | -0.075 | 9.66E-01 | -1.374 | 9.57E-02 |

### The absence of H-NS masks the effect of absence of TsrA at the majority of loci

TsrA (VC0070) has weak amino acid sequence similarity with the N-terminal oligomerization domain of H-NS in *V. cholerae* and was shown to play a role as a transcriptional regulator [44–46]. Because the regulon of TsrA, based on transcriptomic studies, overlaps with that of H-NS [44,45] and because H-NS plays an important role as a global regulator, we tested for epistasis between deletions of *hns* and *tsrA* in the *V. cholerae* clinical isolate under study. Using our RNA-seq results, we assessed the correlation between the regulation of genes from individual deletions of *hns* and *tsrA* (**Figure 4A**). The majority of genes show similar changes in transcript abundance upon loss of H-NS or TsrA (and thus, we hypothesize that H-NS and TsrA co-regulate many of these targets), on the basis of the robust linear regression with a slope close to 1; however, there is a cluster of 49 genes that show substantially more increased transcript levels in the absence of H-NS compared to that of TsrA (thresholding at three standard errors above the curve of slope=1 in the top right quadrant). This cluster includes, for example, the toxin coregulated pilus VC0828/*tcpA* with significant log2FC=6.40/q-value=6.85e-122 upon deletion of *hns*, compared to the effect of *tsrA* deletion with a log2FC=1.79/q-value=3.55e-12.

**Figure 4.**
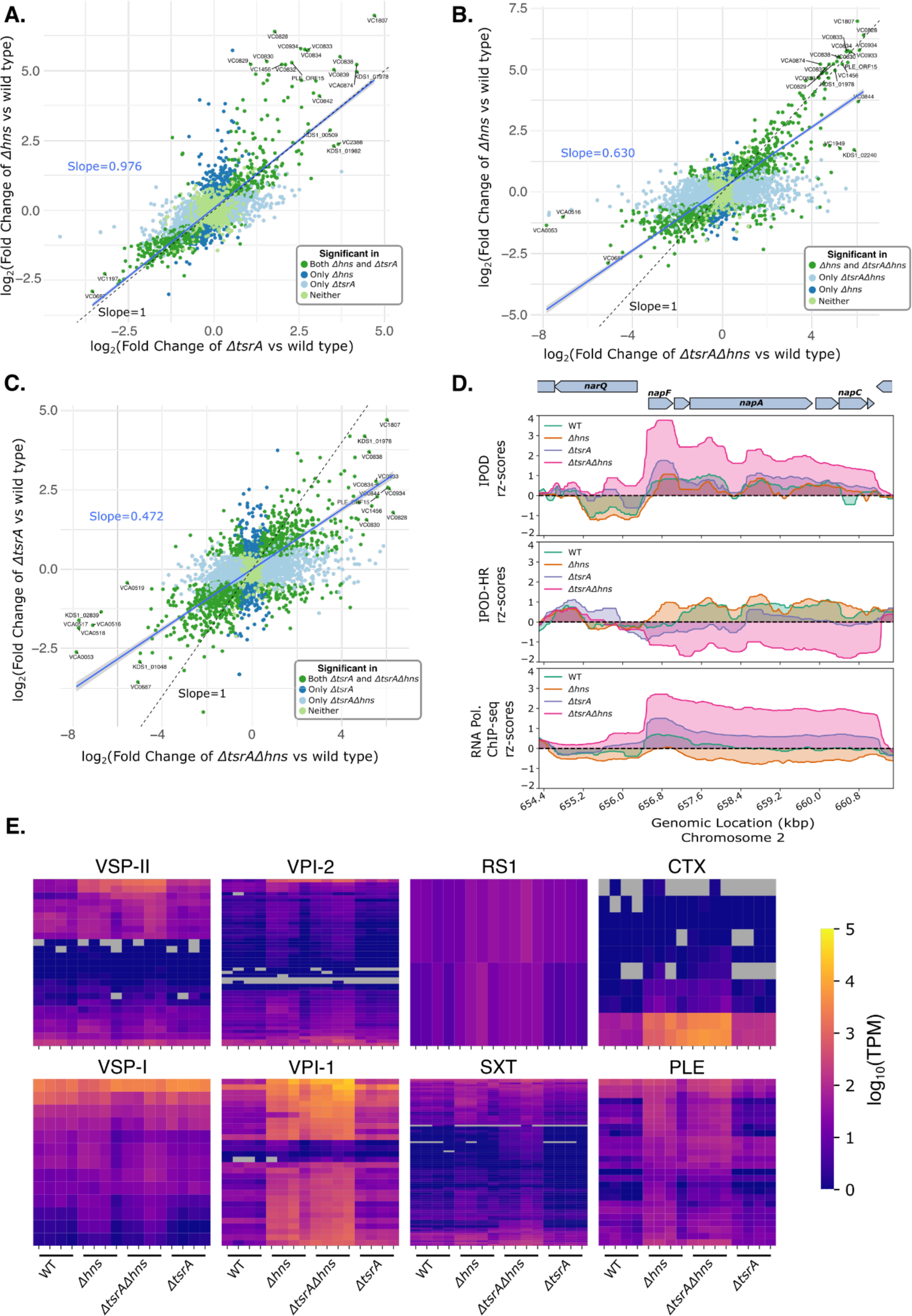
(next page): Effect of the absence of both of H-NS and TsrA on transcript levels in V. cholerae. A-C) Pairwise comparison of log2FC (Fold Changes) in transcript abundances (measured via RNA-seq) for the indicated genotypes, relative to wild type. Blue fitted lines represent the robust linear fit of significantly (q-value less than or equal to 0.1) differentially expressed genes in both of the genotypes (green points) compared; slopes for the fitted lines for each comparison are shown in blue. A line with slope of 1 is shown in black. The points represent four categories with indicated colors: significant genes in both genotypes, significant in only one or the other, significant in neither. The genotypes of pairwise comparisons relative to wild type are: A) *Δhns* vs. *ΔtsrA;* B) *Δhns* vs. *ΔtsrAΔhns;* C) *ΔtsrA* vs. *ΔtsrAΔhns* D) Protein occupancy (total and RNA polymerase) in an EPOD spanning the *napFDABC* locus in cells of the indicated genotypes, during exponential growth in LB media. E) Heatmap representing the transcript abundances (log10[TPMs]) of genes in the strains shown in various HAEs: VSP-I, VSP-II, VPI-1, VPI-2, SXT, RS1, PLE and CTX. The rows represent the genes within the HAEs, the columns represent individual biological replicates of the indicated genotypes. Gray rectangles in heatmaps represent genes with 0 TPMs. Genes in our revised reference genome that mapped to multiple VC numbers were omitted from analysis.

We also compared the effects of individual deletions of *hns* or *tsrA* with those of the double knockout (**Figure 4B,C**). The expression changes caused by the double knockout, especially for genes showing strong differential expression, largely track with the changes seen in the *hns* single knockout (genes along the line of slope=1), again confirming that deletion of *hns* is the primary cause of expression changes in the double knockout. For the HAEs studied here, this pattern of H-NS regulatory dominance largely holds true (**Figure 4E**). However, some exceptions exist: 55 genes are upregulated more strongly in the absence of both proteins out of 381 significantly upregulated genes in *Δhns*, suggesting independent and additive effects of H-NS and TsrA on some loci. For instance, the uncharacterized gene KDS1_02240/s003 [42] (a ParM/StbA family protein based on an NCBI blast search), located in the SXT, shows a large increase in expression in the double knockout (log2FC=5.87/q-value=7.57e-31) compared to the individual deletion strains (log2FC=1.72/q-value=0.0167 in the *Δhns* and log2FC=0.731/q-value=0.253 in the *ΔtsrA*). Similarly, the accessory colonization factor VC0844/AcfA, encoded within VPI-1, is higher in expression (log2FC=6.05/q-value=6.3e-24) in the double mutant compared to individual deletions of *hns* (log2FC=3.69/q-value=1e-6) or *tsrA* (log2FC=2.57/q-value=1e-4), respectively, indicating roughly additive effects of the deletions. These results suggest that both H-NS and TsrA synergistically regulate a subset of genes, but H-NS is responsible for the majority of expression changes in the double knockout. In contrast, at the majority of H-NS regulated sites, the additional loss of TsrA does not have any further impact on regulation, suggesting that TsrA’s role is to act via modulation of H-NS or another DNA-binding protein.

The *napFDABC* (VCA0676 - VCA0680) locus, encoding for periplasmic nitrate reductase [73], provides an example of some exceptions, based on IPOD-HR experiments, where we observe independent regulation by H-NS and TsrA (**Figure 4D**). De-repression of this locus is observed in the absence of TsrA, whereas expression drops in an *hns* knockout; however, the double mutant results in a further increase in RNAP occupancy (and thus presumably transcription) relative to *tsrA* deletion. These results appear to indicate that TsrA and H-NS act independently of each other at the *napFDABC* locus and when they are both no longer present, this chromatin region becomes de-repressed on the basis of increased RNA polymerase occupancy (**Figure 4D**). RNA-sequencing also demonstrates an increase in *napA* transcript level in the double knockout (log2FC=1.28/q-value=6.05e-12) relative to either the *hns* (log2FC=0.61/q-value=1.86e-2) or *tsrA* (log2FC=0.43/q-value=4.76e-2) single mutants. This result suggests that TsrA may act alongside of (and independently of) H-NS at a few loci, similar to the behavior of H-NS paralog StpA in *E. coli,* where we have seen a synergistic de-repression upon the deletion of both *stpA* and *hns [27]*. However, unlike StpA, which binds directly to DNA [26], TsrA has not been predicted to have DNA-binding ability [44–46], suggesting it may function through H-NS and/or another protein.

iPAGE analysis for gene set enrichment is consistent with previous findings that TsrA, like H-NS, regulates many virulence and HAE-associated biological pathways (**Supplementary Figure 6A**). Many metabolism-related GO terms such as “organic acid catabolic process”, “*de novo* IMP biosynthetic process”, and “peptide-transporting ATPase activity” are highly expressed in the absence of both H-NS and TsrA. Consistent with previous studies [44,45], we observe that the regulons of both H-NS and TsrA are both AT-rich, which is a characteristic of many horizontally acquired elements and targets of H-NS in gamma-proteobacteria (**Supplementary Figure 6B**).

To further assess the interplay of regulation by H-NS and TsrA, we investigated the enrichment of GO term pathways that are uniquely upregulated in the single deletion of *hns* or *tsrA* and the double mutant **(Supplementary Figure 7**). We found that the shared regulons of H-NS and TsrA consist primarily of metabolic pathways, including “organic acid catabolic process”, “organic anion transport”, “generation of precursor metabolites and energy”, “tricarboxylic acid cycle enzyme complex” and others. In contrast, the regulatory targets uniquely affected by *hns* deletion and *tsrA* deletion appear to contain disjointed sets of genes involved in host colonization and virulence, including terms such as “host cell plasma membrane” (e.g., VC1451 and VC0822) in *hns* deletion and “pilus” (e.g., VC2423, VC0857 and VC0409) in *tsrA* deletion. These findings indicate that TsrA in fact acts independently of H-NS in regulation of a few key targets involved in virulence, suggesting that TsrA may act with another DNA binding protein to affect gene expression at some loci.

### Structural Modeling Demonstrates a Potential Mechanism for TsrA to Modulate H-NS Binding

As the results above suggest that the effects of TsrA are mediated largely, though not exclusively, through H-NS, we next sought to determine a potential mechanism for this behavior. Consistent with the absence of a recognizable DNA binding domain [44–46,74], modeling a TsrA dimer using DMFold [75] yields a high confidence structure with a predicted quality score (QS; see Materials and Methods) of 0.75, but no clear nucleic acid binding surface is present (**Figure 5A-B**), and a Foldseek [76] search of one of the modeled chains with an E-value cutoff of 1 yields no hits to known DNA-binding proteins. In contrast, modeling of a H-NS/TsrA heterotetramer using DMFold shows a striking potential binding mode in which TsrA inserts itself into an H-NS dimer, losing TsrA-TsrA contacts in favor of a close association with H-NS (**Figure 5C**), with a lower QS of 0.47. We note that the structure generally has high local confidence except in a potentially flexible loop connecting the oligomerization and DNA binding domains of the H-NS. In the modeled heterotetramer, the N-terminal dimerization domain (**Figure 5D**) and the C-terminal DNA binding domain (**Figure 5E**) of H-NS are both individually similar to existing experimental structures of *E. coli* H-NS, but TsrA presents an additional interface directly adjoining the H-NS DNA binding domain (**Figure 5F**). The proximity of TsrA in the heterotetramer model to the DNA binding domain of H-NS in fact presents a unified potential DNA binding interface, providing a ready molecular explanation for how TsrA may modulate H-NS - DNA interactions. While the majority of TsrA effects appear to be H-NS mediated, and explicable through the model shown here, the presence of a minority of H-NS independent TsrA targets (as shown in **Figure 4**) suggests that TsrA may also interact with other regulators besides H-NS, perhaps in some similar mode.

**Figure 5:**
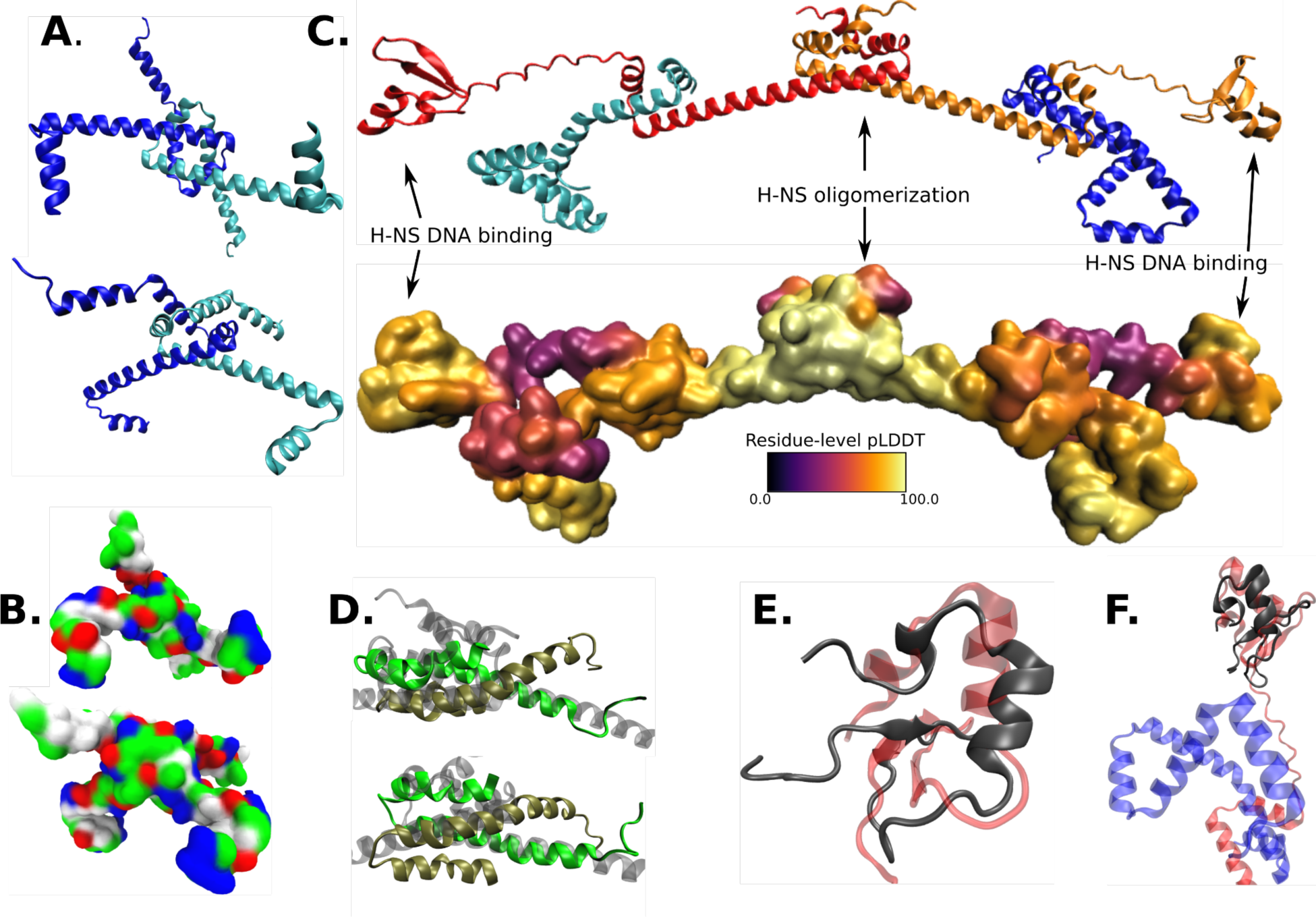
Structural modeling of TsrA/H-NS interactions. A) DMFold model of a TsrA dimer, with the two chains shown in blue and cyan. B) Equivalent view to panel A, showing a surface with residue types colored (white: hydrophobic; green: hydrophilic; blue: basic; red: acidic). C) Model of a 2:2 TsrA/H-NS heterotetramer, with the TsrA chains colored as in panel A, and the H-NS chains colored in red and orange; below is shown a surface representation colored by the residue-level predictive confidence (pLDDT scores). D) Superposition of a crystal structure of the oligomerization domain of *E. coli* H-NS (PDB code 1LR1, shown with chains in green and gold) with the equivalent portion of the heterotetramer model from panel C (transparent gray cartoon). E) Superposition of an NMR structure of the DNA binding domain of *E. coli* H-NS (PDB code 1HNR, in red) with the equivalent region of H-NS from the heterotetramer model in panel C. F) As in panel E, but including the modeled TsrA chain (shown in blue).

### IPOD-HR reveals horizontally acquired elements affected and unaffected by the SOS response

Environmental stresses, such as DNA damage, have been shown to affect the regulation of prophages and antibiotic resistance genes found on HAEs for mobilization by stimulating the SOS response [52,53,77]. Thus, we profiled the genome-wide effect of a DNA damage causing agent, mitomycin C (MMC), on transcription and protein occupancy of *V. cholerae* to identify DNA damage responsive regions of the genome. Transcription and protein occupancy of some of the HAEs were affected by MMC treatment, such as SXT, RS1, CTX, and VSP-1, and many EPODs/nEPODs. (**Figure 6A, B; Supplementary Figure 2, 3, 4**). However, the superintegron and PLE exhibit negative occupancy signal under MMC treatment, which is not associated with changes in RNA polymerase binding, suggesting that H-NS or other factors that are producing negative signal during interphase extraction are perhaps binding these regions to maintain silencing during DNA damage (**Figure 6A-C, Supplementary Figure 3, 4**), or that additional transcriptional activators would be needed to enhance transcription in those regions. SXT, RS1, and CTX show increased RNA polymerase binding compared to untreated cells (**Figure 6D, Supplementary Figure 4,8B**). The SXT integrative conjugative element is a large horizontally acquired genetic element that harbors antibiotic resistance genes and phage defense systems [49,53,78] and it has been shown to be mobilized when the lambda phage cI-like repressor in SXT is proteolyzed by RecA, allowing expression of the genes necessary for the HAE’s transfer that are otherwise repressed [52,53,77]. For example, the *mobI* gene and origin of transfer (*oriT*) of SXT [79], which is located and conserved in the intergenic region of *mobI* and *s003* in many SXT elements (**Figure 6D**), showed increased RNA polymerase binding upon MMC treatment, suggesting increased expression to prepare for mobilization of the element. We also observed a localized region of high protein occupancy in the *mobI-s003* intergenic region, perhaps corresponding to the recruitment of the relaxosome consisting of *oriT* binding proteins needed for the conjugative transfer, as is observed in the bacterial DNA conjugation process [80,81]. Other loci in the SXT exhibiting increased RNA polymerase occupancy during MMC treatment are shown in **Supplementary Figure 8B**, allowing us to assess which regions are activated during SOS response. In contrast to the chromosome 1 HAEs, PLE and the remainder of the superintegron exhibit a strongly negative IPOD signal upon MMC treatment with no concurrent increase in RNA polymerase binding (**Figure 6A-C**). This observation suggests that MMC causes substantial occupancy of PLE by H-NS or some other atypical factor. This occupancy results in a negative signal in our assay, but clearly indicates the presence of a factor with the potential for maintaining repression of these regions.

**Figure 6:**
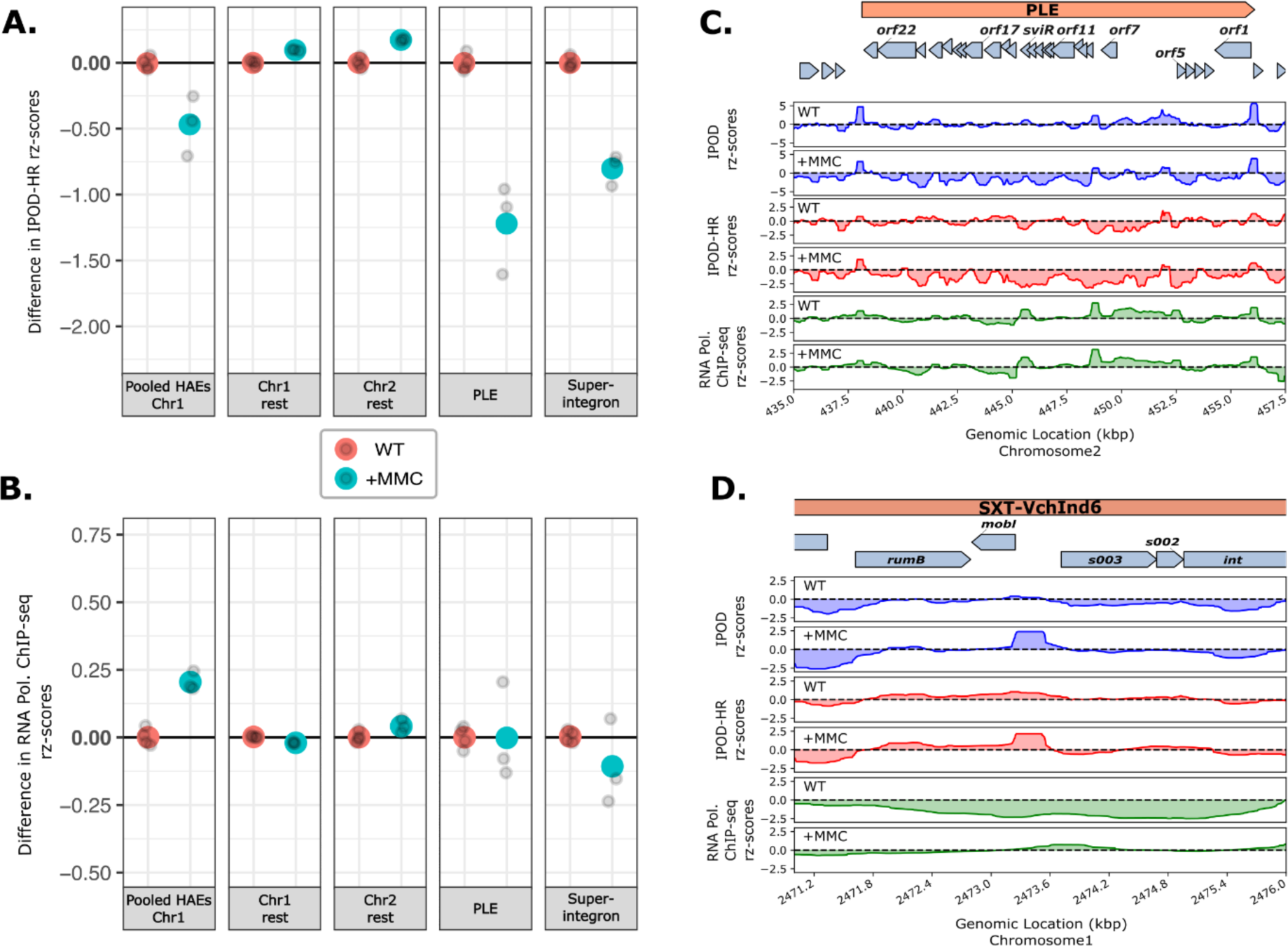
Effect of mitomycin C on the regulation of horizontally acquired elements. A-B) Changes (relative to WT) in robust rz-scores in IPOD-HR (A) and RNA pol occupancy (B) in untreated and mitomycin C treated (+MMC) *V. cholerae* cells over the regions shown. Individual replicates are shown in gray points and the larger colored points are the mean of each of the replicates for each condition. See Fig. 3B for details on how the summary statistics shown here were generated. C) Total protein occupancy signals (IPOD), RNA polymerase ChIP subtracted protein occupancy (IPOD-HR) and the RNA polymerase occupancy (RNAP ChIP-seq) in the PLE island in untreated and MMC treated cells. D) Total protein occupancy signals (IPOD), RNA polymerase ChIP subtracted protein occupancy (IPOD-HR) and the RNA polymerase occupancy (RNAP ChIP-seq) in a region of the SXT element in untreated and MMC treated cells.

## Discussion

Proper regulation of virulence genes and horizontally acquired elements is critically important for bacterial fitness, as constitutive expression would be extremely metabolically costly, but failure to express those genes when needed would prevent optimal host colonization, nutrient acquisition, and growth. Through the application of whole-genome protein occupancy profiling and RNA polymerase ChIP-seq to a series of strains lacking different nucleoid-associated proteins (NAPs), we identified a division of labor across different NAPs for regulating different genetic elements associated with horizontal gene transfer and bacterial virulence. Consistent with previous data, we observe that the NAP H-NS is the main xenogeneic silencer in *V. cholerae* [38,39]; deletion of H-NS was sufficient to trigger increased transcription of many genes. Additionally, deletion of the recently described protein TsrA, a protein with a regulon heavily overlapping that of H-NS, appears to primarily act through H-NS and on H-NS dependent targets, as the regulatory effects of a tandem H-NS/TsrA deletion closely resembled those caused by loss of H-NS alone at the majority of H-NS upregulated targets (381 genes). Because TsrA does not have a predicted DNA-binding domain, it may modulate and associate with H-NS only at certain loci to repress the H-NS targets, much like Hha in *E. coli* [23,82]. However, at some loci (55 in all), deletions of *tsrA* and *hns* show additive or even supra-additive effects, perhaps caused by TsrA associating with another DNA binding protein. Similar observations were made with Hha in *Salmonella* Typhimurium where Hha may have an H-NS-independent effect on gene expression [83]. It is especially notable that some loci regulated by TsrA, even in the absence of H-NS, are involved in host colonization and virulence (e.g., some genes in “pilus” GO term: VC0409, VC0857, VC2423). Future studies directly tracking the binding sites of TsrA and H-NS in each others’ presence and absence, and possibly concerted efforts to identify potential protein-protein interaction partners of TsrA, will be fruitful in more fully resolving the role of TsrA in modulating *V. cholerae* gene expression.

Other abundant NAPs such as integration host factor (IHF) appear to regulate a distinct set of genes from those covered by H-NS. For example, we found that IHF is involved in regulating the overall balance of iron metabolism. Loss of IHF also appears to alter protein occupancy in H-NS regulated regions, perhaps corresponding to competition with H-NS binding **(Supplementary Text)**; functional implications of these changes (e.g., whether the strength of repression by H-NS is increased in the absence of binding from competing factors in HAEs) remains unclear.

Because of the ability of HAEs to mobilize and further promote horizontal gene transfer during DNA damage, we investigated the genome-wide effect of SOS response of *V. cholerae* with IPOD-HR as the induction of SOS response with DNA damaging agent mitomycin C (MMC). We observed that some loci in the HAEs have higher RNA polymerase occupancy in comparison with a no-treatment control. While some regions of HAEs became de-repressed upon the addition of MMC, other HAEs like the phage satellite PLE were not affected in transcription but we observed substantial changes in the protein occupancy profiles in those regions, perhaps suggesting that other factors such as H-NS may be binding and indicating a rearrangement of chromatin structure, but the absence of some essential signal, presumably ICP1-encoded, that would be needed for full activation of the HAE.

The H-NS dependent silencing of PLE revealed here appears in stark contrast to the regulatory strategies for HAEs similar to PLEs that rely on helper phages. For example, for the well-studied *Staphylococcus aureu*s pathogenicity islands (SaPIs) – the HAE itself encodes a master repressor (in this case, Stl) that keeps the SaPI in the prophage-like state [84]. Stl-mediated repression is counteracted by Stl complexing with specific phage antirepressors that disrupt the formation of Stl-DNA complex [85,86]. Based on our data, we hypothesize that H-NS is silencing the majority of the PLE and thus not relying on a PLE-encoded master repressor, but still requiring a phage-encoded protein to relieve the repression by H-NS and TsrA (possibly through induction of some transcriptional activator that remains to be identified). Some examples of phage proteins that relieve the repression by an H-NS family protein (MvaT) include the *Pseudomonas* phage LUZ24 Gp4, which is proposed to inhibit MvaT-DNA bridged complex formation [87]. Thus, similar investigations may lead to the identification of the ICP1-encoded factor(s) needed for full activation of the PLE during ICP1 infection.

We expect that future experiments assessing the extent to which DNA damage or NAP deletions license transcription of the PLE and similar regions will be highly informative (e.g., by testing whether the induction kinetics of PLE are altered by the changes in chromatin structure that we observed), as will additional efforts to determine the proximal signal (presumably some element from ICP1) triggering PLE activation during phage infection.

## Acknowledgments

This work was supported by NIH R35 GM128637 (to PLF) from the National Institute of General Medical Sciences and 1R01AI127652 (to KDS) from the National Institute of Allergy and Infectious Diseases; its contents are solely the responsibility of the authors and do not necessarily represent the official views of the National Institute of Allergy and Infectious Diseases, the National Institute of General Medical Sciences, or NIH. KDS holds an Investigators in the Pathogenesis of Infectious Disease Award from the Burroughs Wellcome Fund. DTD was supported by the National Science Foundation Graduate Research Fellowship (2018257700). The authors are grateful to Dr. Jeremy Schroeder for developing, and assisting with, the IPOD-HR data analysis pipeline, and to Dr. Rebecca Hurto for technical assistance.

## Data availability

IPOD-HR and RNA-sequencing data are available on GEO (accession number: GSE250408) with the URL:

https://www.ncbi.nlm.nih.gov/geo/query/acc.cgi?acc=GSE250408. Nanopore sequencing data for KDS1 are available from the Sequencing Read Archive (SRA) with accession PRJNA1056466.

## Materials and methods

### Strains, culture, and media conditions

The genotypes of all *V. cholerae* strains used in this study are listed in Table S2. *V. cholerae* KDS1, an El tor clinical isolate that encodes PLE1, was used as the wild type strain for all experiments and genetic manipulation. All standard overnight bacterial cultures were grown from a single colony in 2 mL volume with aeration at 37°C in standard LB Miller liquid growth medium, supplemented with streptomycin (100 µg/mL).

For cultures requiring mitomycin C treatment, an overnight culture grown in standard conditions was diluted to an OD600 of 0.003 into 140 mL of pre-warmed LB Miller liquid medium supplemented with mitomycin C to a final concentration of 20 ng/mL. Cultures were grown to a final OD600 of 0.3 as described and processed according to the methods below.

### Strain engineering

All genetically manipulated strains were generated through splicing by overlap extension (SOE) PCR to generate PCR products with FRT-Spec^R^/Kan^R^-FRT in place of the gene of interest. This SOE PCR product was added to *V. cholerae* grown overnight on chitin to induce natural competence and transformants were selected with the appropriate antibiotics, as previously described [88]. Transformants were screened by colony PCR and Sanger sequencing to confirm the presence of the deletions of interest.

### IPOD sample preparation

IPOD samples were collected as previously described [51] with minor modifications. Overnight cultures were diluted to a starting OD600 of approximately 0.003 in a final volume of 140 mL of LB medium in a 1 L flask. The diluted cultures were grown at 37°C with shaking at 250 RPM until a final OD600 of 0.3. 28.9 mL of culture was collected at each time point and mixed with 300 μL 1 M sodium phosphate buffer (pH 7.4) and 810 μL formaldehyde (37%, fresh) in a 50 mL conical tube, then allowed to crosslink for 5 minutes shaking at room temperature (notably, unlike in [51], no rifampicin was added prior to crosslinking). After 5 minutes, the crosslinking reactions were quenched with 6 mL of 2M glycine and returned to the room temperature shaker. After 10 minutes of shaking, tubes were placed in ice for 10 minutes and then pelleted at 7,000x g at 4°C for 4 minutes. Pellets were washed twice with 10 mL of ice-cold phosphate-buffered saline and the final cell pellets were snap-frozen in a dry ice-ethanol slurry and stored at −80°C.

### Cell lysis, digestion, and lysate clarification

The original IPOD-HR protocol is from ref. [51]; below we summarize the procedures applied here. Frozen cell pellets were resuspended in 600 μL of 1X IPOD lysis buffer (10 mM Tris HCl, pH 8.0; 50 mM NaCl) with 1x protease inhibitors tablet (Roche Complete Mini, EDTA free, Roche Diagnostics GmbH, Mannheim, Germany) and 1.5 μL lysozyme (ThermoScientific, REF90082, 50 mg/mL) incubated for 15 minutes at 30°C with gentle shaking and then placed on ice. Resuspended cells were sonicated with a Branson sonicator at 25% power with three bursts of 10s ON and 10s OFF while in an ice water bath. Sonicated cells were then digested with 6μL RNaseA (SIG 10109169001, Sigma-Aldrich/Roche, 10mg/mL), 5.4 μL of 100mM MnCl2, 4.5 μL of 100mM CaCl2, 9 μL of DNase I, RNAse-free (ThermoScientific, #89836, 1U/μL) and incubated on ice for 30 minutes, after which reactions were quenched with 50 μL 500mM EDTA (pH 8.0); we aimed to obtain fragment sizes of about 150 bp. 400 μL of 1x IPOD lysis buffer without protease inhibitors and lysozyme was added to the digested mixture and vortexed. The mixture was clarified by centrifugation for 10 minutes at 15,700 x g at 4°C and transferred to new tubes. The clarified lysate was partitioned into three different tubes: 50 μL for INPUT (for baseline reference), 400 μL for IPOD (for total protein occupancy), and the rest for CHIP (for RNA polymerase occupancy). 450 μL of ChIP elution buffer (50mM Tris (pH 8.0), 10mM EDTA, 1% SDS) was added to the INPUT sample and kept on ice until reverse cross-linking step (mentioned in the next step).

### Interphase extraction and nucleic acid purification

400 μL of 100 mM Tris Base and 800 μL of 25:24:1 phenol:chloroform:isoamyl alcohol (Sigma Aldrich, #77617) was added to the 400 μL of clarified lysate kept for the IPOD-HR sample, vortexed for 10s, and then incubated for 10 minutes at room temperature. To obtain a separation of the organic and aqueous layers and formation of the white interphase disc, which is enriched in protein-DNA complexes, the mixture was spun for 2 minutes at 21,130 x g at room temperature. The aqueous and organic layers around the interphase were removed while minimizing disturbance of the interphase disc. The extracted disc was washed once in 350 μL TE (10 mM Tris, pH 8.0; 1 mM EDTA), 350 μL 100 mM Tris base, and 700 μL 24:1 chloroform:isoamyl alcohol. The layers were again separated by spinning for 2 minutes at 21,130 x g at room temperature. After removal of liquid around the interphase disc, a final wash of 700 μL TE and 700 μL 24:1 chloroform:isoamyl alcohol was applied. After vortexing and spinning the final wash as before, as much of the liquid around the white disc as possible was removed and discarded. The interphase disc was resuspended in 500 μL of ChIP elution buffer and vortexed. The resuspended interphase of IPOD sample and INPUT sample (kept on ice from above step) were incubated at 65°C overnight to reverse-crosslink (6-16 hours).

### RNA Polymerase Chromatin Immunoprecipitation

The rest of the clarified lysate (about 500 μL) was mixed with 1 volume of 2x immunoprecipitation (IP) buffer (200mM Tris (pH 8.0), 600mM NaCl, 4% Triton X-100) and incubated with 10μg of anti-*E. coli* RNA polymerase antibody (NeoClone WP023, NeoClone, Madison, WI, Lot: 2019G15-002) overnight with rocking at 4°C. After overnight incubation, 50 μL of protein G beads (New England Biolabs (NEB), S1430S) per sample was equilibrated in 1x IP buffer. 50 μL of equilibrated protein G beads was added to each antibody-lysate mixture and then incubated with rocking 2 hours at 4°C. After incubation, the beads were washed once in 1mL of the following buffers: IP wash buffer A, IP wash buffer B, IP wash buffer C, 1x IP buffer, and 1x TE.

- 1x wash buffer A (100mM Tris, pH 8.0; 250mM LiCl; 2% Triton X-100; 1mM EDTA)
- 1x wash buffer B (10mM Tris, pH 8.0; 500mM NaCl; 1% Triton X-100; 0.1% sodium deoxycholate; 1mM EDTA)
- 1x wash buffer C (10mM Tris, pH 8.0; 500mM NaCl; 1% Triton X-100; 1mM EDTA)

After the wash steps, beads with the immunoprecipitated material were resuspended in 500 μL of ChIP elution buffer by rocking for 5 minutes; the samples were then incubated for 30 minutes at 65°C with vortexing every 5 minutes. The beads were then separated and discarded to retain only the eluted immunoprecipitated sample. The samples were then left to reverse-crosslink overnight (6-16 hours) at 65°C.

### DNA extraction post reverse cross-linking

The DNA from INPUT, IPOD, and ChIP samples were isolated and purified following identical steps. After overnight incubation to reverse the formaldehyde crosslinks, each sample was incubated with 10 μL of RNAse A (Roche Diagnostics GmbH, #SIG10109169001, 10mg/mL) for 2 hours at 37 °C. Then we followed by adding 10 μL Proteinase K (ThermoScientific, #EO0491, 20mg/mL) and incubating for 2 hours at 50 °C. The DNA of each sample was isolated by phenol-chloroform extraction using one volume of 25:24:1 phenol:chloroform:isoamyl alcohol, then re-extracted with one volume of 24:1 chloroform:isoamyl alcohol. At the last stage the samples went into DNA LoBind tubes. The isolated DNA was then precipitated by adding 1/25^th^ volume of 5M NaCl as a precipitating salt, 1/300^th^ volume of GlycoBlue (Invitrogen, #AM9515, 15mg/mL) as a co-precipitant, and two volumes of ice cold 1:1 isopropanol:ethanol. The DNA was precipitated at 4°C for one hour and at −20 °C for more than 1 hour or overnight.

Chilled samples were then centrifuged for 15 minutes at 16,100 x g at 4 °C to pellet the DNA. All the liquid was then removed without disturbing the DNA pellet. Pellets were then washed in freshly diluted 95% ethanol, vortexed and then centrifuged for 5 minutes at 16,100 x g at 4°C. Finally, the liquid was removed, and the remaining ethanol was left to evaporate for 30 minutes or less. The following volumes of TEe (10 mM Tris, pH 8.0; 0.1 mM EDTA) were added to resuspend the DNA from the three sample types: 200 μL for INPUT pellets, 50 μL for IPOD, and 30 μL for ChIP. All samples were quantified with QuantiFluor dsDNA System (Promega, #2670) using a BioTek Synergy plate reader. Input samples were assessed for fragment sizes on 2% agarose gel electrophoresis.

### Illumina library preparation for IPOD-HR

The purified and quantified DNA from INPUT, IPOD, and RNA Polymerase ChIP samples were then prepared for Illumina sequencing using NEBNext^®^ Ultra™ II DNA Library Prep Kit for Illumina^®^ (#7645S/L) with NEBNext^®^ Multiplex Oligos for Illumina^®^ Unique Dual Index UMI Adaptors DNA Set 1 (#E7395S) or NEBNext^®^ Multiplex Oligos for Illumina^®^ Dual Index Primers Set 1 (#E7600S) and Dual Index Primers Set 2 (#E7780S). Omega Biotek Mag-Bind DNA purification or Axygen purification beads were used for all SPRI bead cleanup steps. The bead purification at the post-adaptor ligation stage was modified as follows: 1.8x volume of DNA purification beads and 0.7x isopropanol were added instead of the normal 0.9x volume of beads.

Each sample prepared for sequencing was quantified and assessed for fragment size and for the presence of any adaptor present on 2% agarose gel electrophoresis. The samples were pooled into libraries, which were sequenced on a NextSeq 550 instrument at the Michigan Advanced Genomics Core (GEO accession: GSE250408).

### RNA isolation

Bacterial strains of interest were grown to an OD600 of 0.3 in 4 mL culture tubes and mixed 1:1 with ice-cold methanol in a 15 mL falcon tube, under culture conditions equivalent to the IPOD experiments described above. Methanol-treated samples were pelleted at 7,000 x g at 4°C, and the resulting pellets were washed in ice-cold phosphate-buffered saline. The washed pellets were resuspended in 200 µL TRI Reagent (Millipore/Sigma) and incubated for 5 minutes at room temperature. The samples were mixed with 40 µL chloroform, vortexed, and incubated for 10 minutes at room temperature. The chloroform-treated samples were then centrifuged at 12,000 x g for 10 min at 4°C. Following centrifugation, the upper (aqueous) phase was collected, promptly mixed with 110 µL 2-propanol and 11 µL pH 6.2 3M sodium acetate, and mixed vigorously. The samples were centrifuged at 12,000 x g for 15 min at 4°C, and the pellets were washed twice with 500 µL of 75% ethanol. The washed pellets were incubated uncapped at 65°C for 3 minutes to evaporate any residual ethanol and resuspended in 20 µL of diethyl dicarbonate (DEPC) treated water.

### rRNA depletion and RNA-sequencing library preparation

Four replicates of WT, *ΔtsrAΔhns, ΔtsrA,* and *Δhns,* and three replicates of *ΔihfA* of purified RNA were used. Each purified RNA sample was subjected to Baseline-ZERO™ DNase digestion in a 100 μL reaction of 20 μL of purified RNA, 5 μL Baseline-ZERO™ DNAse enzyme (LGC Biosearch Technologies, Pt# E0110-D1, 1 U/μL), 10 μL 10X Baseline-ZERO™ DNase Reaction Buffer, 2 μL RNase Inhibitor, Murine (NEB, #M0314S/L) and nuclease-free water at 37°C for 30 minutes. The RNA was purified with Zymo RNA clean and concentrate kit and eluted with nuclease-free water. The quality of purified RNA was assessed on a 2% agarose gel with guanidine thiocyanate and quantified with a NanoDrop spectrophotometer. Purified RNA was depleted of ribosomal RNA (rRNA) using a NEBNext^®^ rRNA Depletion Kit (Bacteria) (NEB, #E7850L/X) and library preparation was done using NEBNext^®^ Ultra^™^ II Directional RNA Library Prep Kit for Illumina^®^ (NEB, #E7760S/L), according to the manufacturer’s instructions. The libraries were assessed for quality with agarose gel electrophoresis and quantified with a Promega Quantifluor High Sensitivity dsDNA kit.

### IPOD-HR pipeline

Raw read data from the IPOD, ChIP and INPUT samples were demultiplexed using bcl2fastq2 software (Illumina). The data were then processed using version 2.7.2 of the IPOD-HR pipeline via a singularity container (accessible at https://github.com/freddolino-lab/ipod). The peak calling was performed with version 2.8.1. Briefly, sequences are processed to remove the adapters with cutadapt (4.3 with Python 3.8.6) [89] and low-quality reads were trimmed with trimmomatic (version 0.39) [90], then the processed reads were aligned with bowtie2 (version 2.4.4) [91] to the assembled reference genome of KDS1 strain of *V. cholerae* in this study. The quality of reads and alignment were assessed with chipqc and fastqc. The reads were quantile normalized and log ratios of IPOD vs INPUT and ChIP-seq vs INPUT were calculated. Further, to obtain occupancy scores without RNA polymerase bound regions, the pipeline subtracts RNA polymerase ChIP-seq normalized scores from the IPOD vs INPUT normalized scores. The pipeline generates bedgraph and narrowpeak files with the normalized scores and we use them for the analysis of data with in-house R, shell and python scripts as described throughout the text.

### Permutation Test for EPODs and nEPODs

A permutation test was performed to identify whether the AT percentage in called EPODs or nEPODs was significantly different from the background (i.e., the remainder of the genome excluding EPODs and nEPODs). We applied the same method separately to EPODs and nEPODs. First, we obtained 1000 randomized EPOD (or nEPOD) distributions containing EPODs of the same length and total number as our real EPODs with bedtools (version 2.30.0) [92], making sure that randomized EPODs excluded the nEPODs, to generate the null distribution of EPODs. For each randomized set of EPOD locations, we then calculated the difference in AT content between the shuffled EPOD locations and the corresponding background, thus obtaining a summary statistic sampled from the null distribution. We compared these values to the difference in AT percentage median between the real EPODs and the real background. The p-value was then calculated as follows: p-value = (# permutations greater than real EPODs + 1)/(1000 permutations + 1)

### Genomic reference sequence and annotations

We obtained an initial genome reference sequence for KDS1 by sequencing high molecular weight genomic DNA purified from the strain. Raw reads are available at the SRA in PRJNA1056466. Initial assembly was performed on the Nanopore reads (Nanopore sequencing performed at SeqCenter in Pittsburgh, Pennsylvania) using NECAT (version 0.0.1 20200803) [93], resulting in two contigs which clearly correspond to the two *V. cholerae* chromosomes. The assembly was then polished using pilon (version 1.24) [94] using default arguments, with all available reads from INPUT samples (aligned using bowtie2 [91]) as the short read inputs. Manual finishing was performed using a set of iterative rounds of breseq (version 0.37.0) [95] runs to identify remaining discrepancies between the Illumina short read data and the in-progress assembly and resolving them by applying differences with gdtools [95]. We then manually assigned the starting position of each chromosome to match those of commonly used El Tor reference genomes.

Annotation of the newly obtained assembly was performed using prokka (version 1.14.5) [96] with arguments --genus Vibrio --species cholerae --accver 2, using a draft genome obtained from the ref. [97] as a reference for potential proteins. We then assigned VC numbers to all identifiable genes matching the reference El Tor strain (EMBL reference sequences AE003852.1 and AE003853.1 for chromosomes 1 and 2, respectively). Annotated genes from the El Tor reference were aligned to those of our prokka-annotated KDS1 assembly, using nucmer (v. 3.1 [98]) with default settings. We further required that for each potential match, the starting and ending positions of the reference El Tor version and the KDS1 version in the identified features differed by no more than 20 nucleotides.

### RNA-sequencing analysis

RNA sequencing raw read data samples were demultiplexed analogously to the above IPOD-HR pipeline. The Illumina adapter sequence from the raw reads were cut with cutadapt (version 4.1 with Python 3.8.6) [89]: cutadapt -j 24 --quality-base=33 -a AGATCGGAAGAGCACACGTCTGAACTCCAGTCA -A AGATCGGAAGAGCGTCGTGTAGGGAAAGAGTGT

Low quality reads were trimmed with trimmomatic (version 0.39) [90]:

trimmomatic PE -threads 24 -phred33 -validatePairs

TRAILING:3 SLIDINGWINDOW:4:15 MINLEN:14.

The preprocessed reads were pseudoaligned to the transcriptome of KDS1 which was obtained based on our genome assembly/annotation (see above) and quantified with Kallisto (version 0.48.0) [99].

Arguments for indexing: -k 21 –make-unique

Arguments for quantitation: -t 4 -b 200 –rf-stranded –bias

Differential expression calling of pseudoaligned and quantified RNA-seq reads was performed with the Sleuth R package (version 0.30.1) [100] using Sleuth response error measurement (full) model where we fitted our model by condition ∼ batch parameter where condition refers to each genotype: wild type, *ΔtsrAΔhns, ΔtsrA, Δhns,* and *ΔihfA*. The Wald test was performed to obtain differential expression values between the condition parameters and the wild type. Downstream data analysis was performed with in-house R and python scripts. The Gene Ontology (GO) terms utilized for the purple density plot above the volcano plot of *ΔihfA* vs wild type (Figure 2B) were: GO:0006826 GO:0006879 GO:0006880 GO:0010039 GO:0010106 GO:0033212 GO:0033214 GO:0034755 GO:0034756 GO:0034757 GO:0055072 GO:0071281 GO:0097577 GO:0098706 GO:0098711 GO:1901678 GO:0005381 GO:0005506 GO:0015093 GO:0015603, excluding the ChIP-seq-identified Fur regulon genes from ref. [67]. The numbers of genes (55 out of 381) identified as strongly affected in the *ΔtsrAΔhns* were filtered to only include the genes with log2FC above 3 times the mean of standard error in the log2FC of *Δhns* (0.9138129) after filtering the genes VC0070(*tsrA*) and VC1130(*hns*) that were deleted.

### Mapping of H-NS ChIP-seq from *V. cholerae* C6706 to KDS1 strain

The SRA read files from H-NS-V5 ChIP-seq and input control from the published study [71] were converted to fastq.gz files with the SRA toolkit (https://github.com/ncbi/sra-tools). The fastq files were then preprocessed to clip the adaptors with cutadapt (4.1 with Python 3.8.6) [89]:

cutadapt -j 24 --quality-base=33 -a AGATCGGAAGAGCACACGTCTGAACTCCAGTCA -A AGATCGGAAGAGCGTCGTGTAGGGAAAGAGTGT

We then aligned the preprocessed fastq.gz files to the KDS1 reference genome using bowtie2 (version 2.4.4) [91]:

bowtie2 -x reference_index -U fwd_reads.fq.gz -S sam_out -q –local –very-sensitive -p 24 –no-unal –phred33

The sam files were converted to sorted bam files with samtools (version 1.9, using htslib 1.9) [101]. The sorted bam files were quantified using bedtools (version 2.30.0) [92] to obtain aligned and quantified alignment bedgraph files.

The quantified bedgraph files were normalized by rescaling the occupancy data so that the position-wise trimmed mean of each track was 100 (after excluding the top and bottom 5% of the positions), and then a pseudocount of 1 was added to each position. The extracted and input samples were each averaged across replicates for data from each experiment; final occupancy traces for the analysis displayed were the log ratios of each extracted sample relative to the corresponding input.

### iPAGE GO term enrichment analysis

iPAGE analysis was performed with version 1.2a. We used iPAGE in discrete mode separately for EPODs and nEPODs, where the input file had two columns: the location, and a number either 1 or 0, where EPODs or nEPODs were assigned to bin 1 and the background was assigned into bin 0. The gene annotation sets were derived by merging current annotations from the Uniprot El Tor proteome (taxon ID 243277; assigned to genes in our new genome following the procedure described above) with those resulting from running ATGO [102] on the called ORFs in our newly derived reference genome. The iPAGE non-default command line arguments were: --exptype=discrete For RNA-sequencing, we used the same iPAGE version but in a continuous mode, where we assigned 7 total bins to separate the GO term classification. The input files for each strain comparison had two columns: gene label and the wald statistic which is the b value divided by se_b from Sleuth RNA-seq analysis. Command line arguments were: --exptype=continuous --ebins=7 --max_p=0.05

The iPAGE on Supplementary Figure 7 was run in discrete mode, where the input file had the following two columns: genes and discrete categories 0 (all of the rest of the genes that were not upregulated in none of the three categories above 3 standard errors), 1 (upregulated in Δ*hns*Δ*tsrA* that exclude the genes upregulated in Δ*hns*), 2 (uniquely upregulated in Δ*hns*), and 3 (uniquely upregulated in Δ*tsrA*). The iPAGE non-default command line arguments were: --exptype=discrete

