## Supplementary Text and Figures/Tables for "Nucleoid-associated proteins shape the global protein occupancy and transcriptional landscape of a clinical isolate of *Vibrio cholerae*"

### H-NS is likely causing negative occupancy signal

While there is no large change in RNA polymerase occupancy in the *ΔtsrA,* and *Δihf* strains in the VPI-1 **(Figure 3C)**, the occupancy scores (IPOD and IPOD-HR rz-scores) are more negative for these strains compared to the *ΔvctA* control gene and the wild type strain. Although no large change in RNA polymerase occupancy is observed, *ΔtsrA* and *ΔihfA* strains still resulted in a more negative IPOD and IPOD-HR signal in this HAE compared to the wild type or the control gene deletion of VctA. This motivated us to assess the distribution of occupancy scores in the H-NS bound regions from a published study of V5-tagged H-NS ChIP-seq [71] from the strain of *V. cholerae* C6706 that we re-mapped to our wild type KDS1 reference genome **(Supplementary Figure 5C)**. The occupancy signals before and after RNA polymerase subtraction based on IPOD and IPOD-HR for the wild type strain are negative suggesting that H-NS may be a protein in our method where it produces negative signal due to its depletion from the interphase as was the case with Rok in *B. subtilis [21]*. We further assessed the distribution of occupancy scores at these regions in the *ΔihfA* and observe increased distribution of negative scores in the H-NS-bound regions from C6706, suggesting that the absence of IHF results, perhaps, in more H-NS binding, as it was demonstrated for the *tcpA* promoter in the absence of *ihfA* from cells grown in stationary phase in tryptic soy broth [40], or it allows the signal of H-NS occupied regions to be more negative due to lack of competition from other factors binding to the same regions. With this in mind we suggest that regions producing the negative occupancy signal may be due to H-NS.

This is again consistent with the interpretation that *V. cholerae* H-NS yields a negative IPOD signal due to depletion from the interface under the conditions used here; thus, lack of competition from other NAPs leads to an increased negative occupancy signal at H-NS bound regions due to reduced competition (either for binding or for interface partitioning), whereas deletion of H-NS leads to increased transcription of the same regions. However, we do not see large changes in RNA polymerase occupancy in the individual deletions of *tsrA* or *ihfA*, corroborating again that H-NS is still largely repressing CTX region **(Figure 3D, Supplementary Figure 4, 5B)**. In future studies, it will be important to demonstrate what constitutes the negative occupancy during IPOD-HR in *V. cholerae*, whether it is mainly H-NS or a mixture of proteins and why the negative occupancy signal appears strongly when proteins known to act at those loci such as TsrA and IHF are absent. A similar question remains unanswered for the SOS-induced effect on protein occupancy, where some regions produce more negative occupancy signals than others, such as the PLE **(Figure 5C)**, perhaps suggesting the binding of H-NS or other protein factors to this region during DNA damage.

**Supplementary Material**

Supplementary Figure1:

A) GO term enrichment classification analysis of RNA-sequencing results of indicated genotypes: Δ*ihfA*

B) AT percentage of differentially expressed genes in Δ*ihfA* versus wild type. Significant differentially expressed genes (q-value less than or equal to 0.1 are indicated in black and q-value of greater than 0.1 are indicated in gray).

A.

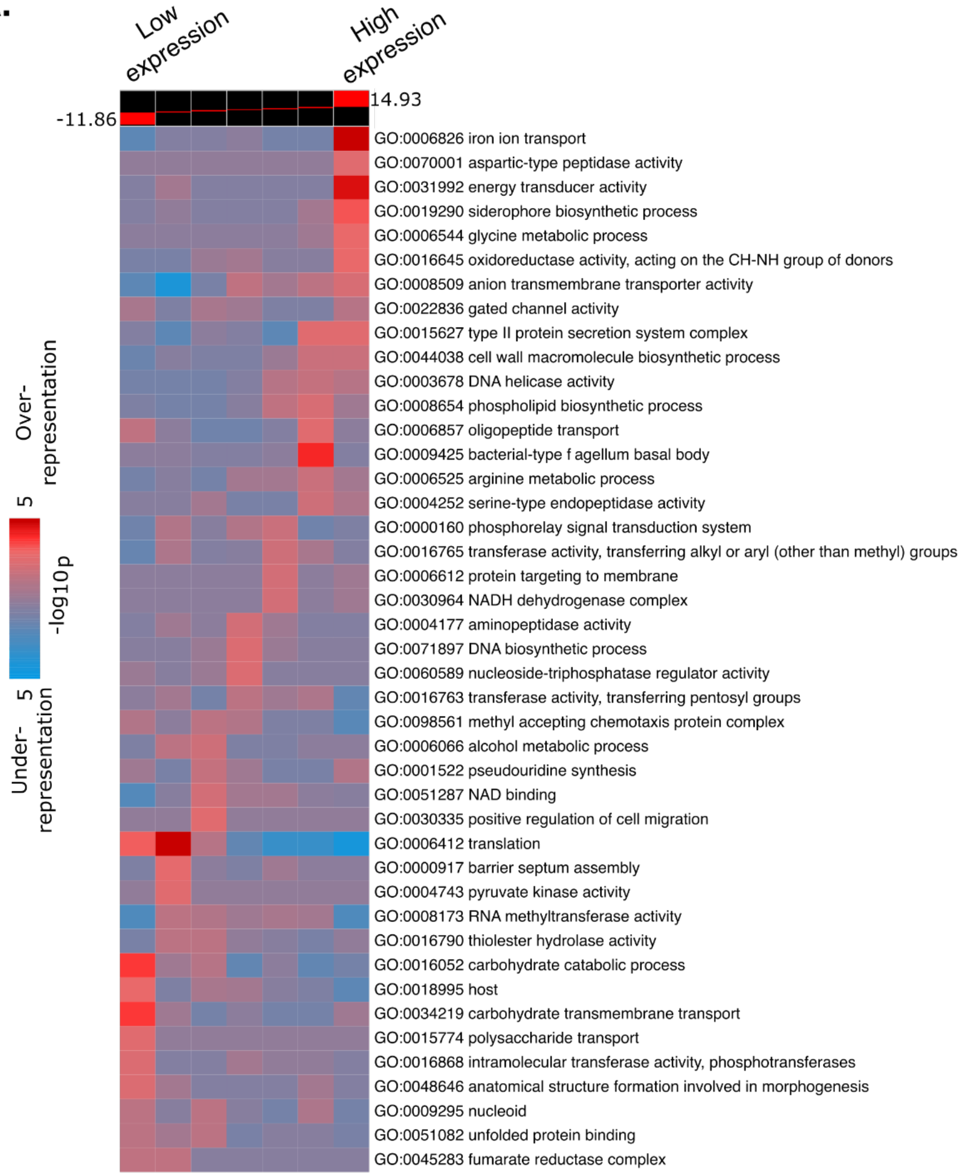
Low expression
High expression
14.93
-11.86
Over-representation
5
-log10p
5
Under-representation
GO:0006826 iron ion transport
GO:0070001 aspartic-type peptidase activity
GO:0031992 energy transducer activity
GO:0019290 siderophore biosynthetic process
GO:0006544 glycine metabolic process
GO:0016645 oxidoreductase activity, acting on the CH-NH group of donors
GO:0008509 anion transmembrane transporter activity
GO:0022836 gated channel activity
GO:0015627 type II protein secretion system complex
GO:0044038 cell wall macromolecule biosynthetic process
GO:0003678 DNA helicase activity
GO:0008654 phospholipid biosynthetic process
GO:0006857 oligopeptide transport
GO:0009425 bacterial-type f agellum basal body
GO:0006525 arginine metabolic process
GO:0004252 serine-type endopeptidase activity
GO:0000160 phosphorelay signal transduction system
GO:0016765 transferase activity, transferring alkyl or aryl (other than methyl) groups
GO:0006612 protein targeting to membrane
GO:0030964 NADH dehydrogenase complex
GO:0004177 aminopeptidase activity
GO:0071897 DNA biosynthetic process
GO:0060589 nucleoside-triphosphatase regulator activity
GO:0016763 transferase activity, transferring pentosyl groups
GO:0098561 methyl accepting chemotaxis protein complex
GO:0006066 alcohol metabolic process
GO:0001522 pseudouridine synthesis
GO:0051287 NAD binding
GO:0030335 positive regulation of cell migration
GO:0006412 translation
GO:0000917 barrier septum assembly
GO:0004743 pyruvate kinase activity
GO:0008173 RNA methyltransferase activity
GO:0016790 thiolester hydrolase activity
GO:0016052 carbohydrate catabolic process
GO:0018995 host
GO:0034219 carbohydrate transmembrane transport
GO:0015774 polysaccharide transport
GO:0016868 intramolecular transferase activity, phosphotransferases
GO:0048646 anatomical structure formation involved in morphogenesis
GO:0009295 nucleoid
GO:0051082 unfolded protein binding
GO:0045283 fumarate reductase complex

B.

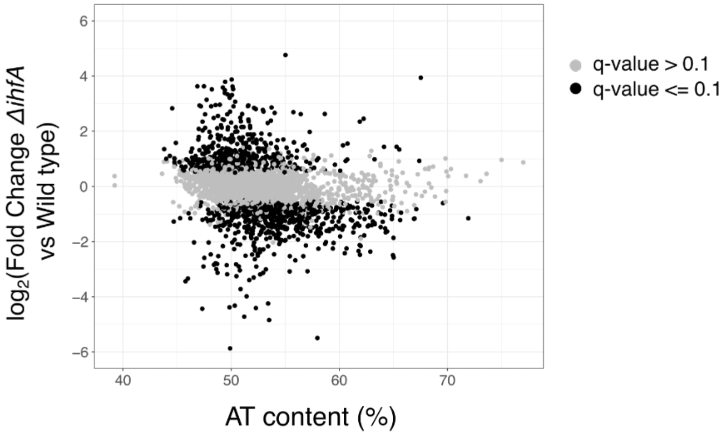
log2(Fold Change ΔihfA vs Wild type)
6
4
2
0
−2
−4
−6
40
50
60
70
AT content (%)
q-value > 0.1
q-value <= 0.1

Supplementary Figure 2:

A) Violin plots of mean differences in EPODs between the average of indicated genotypes and the wild type. Facet grids represent IPOD, IPOD-HR and RNA polymerase ChIP-Seq.
B) Violin plots of mean differences in negative EPODs (nEPODs) between the average of indicated genotypes and the wild type. Facet grids represent IPOD, IPOD-HR and RNA pol ChIP-Seq.

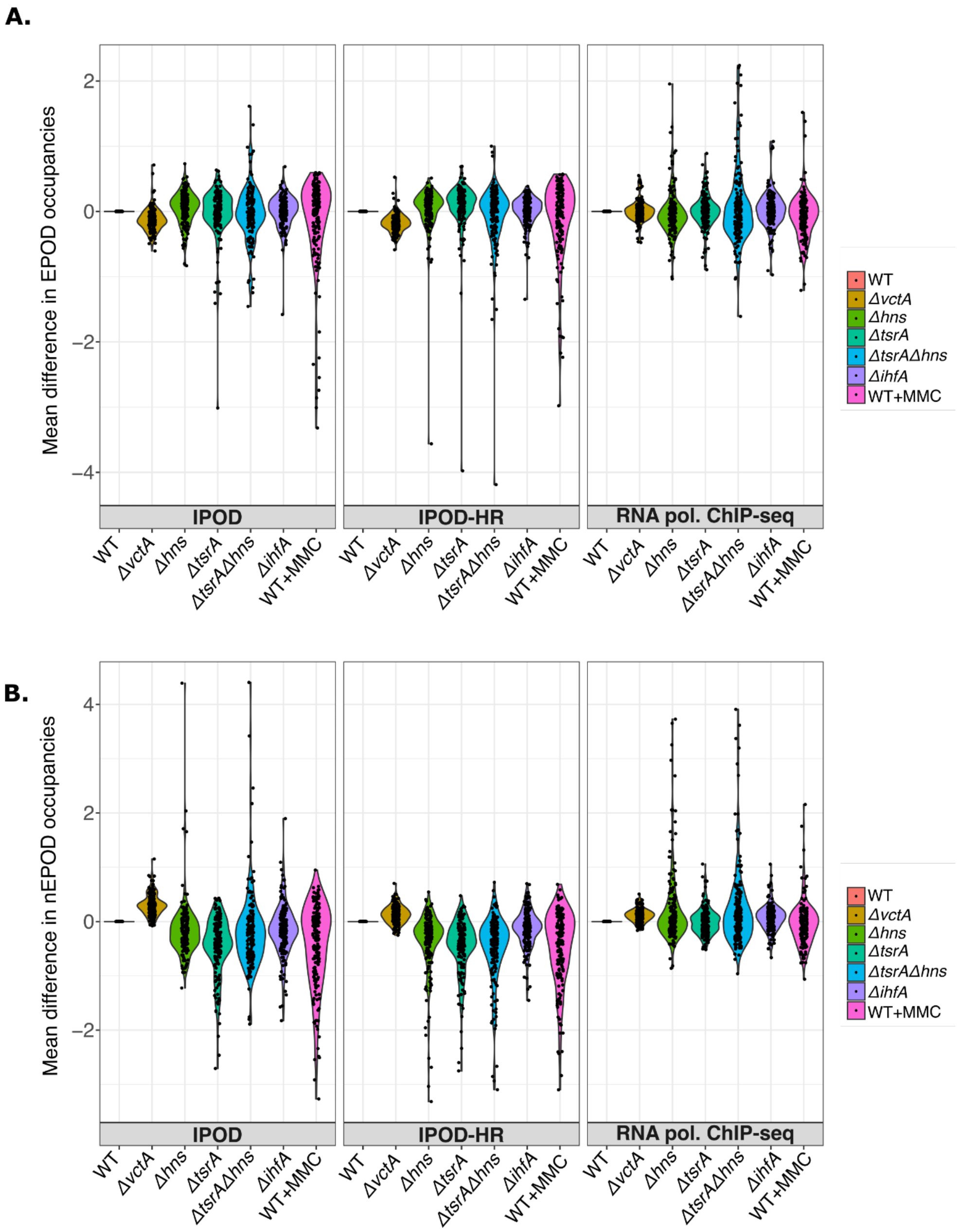

Supplementary Figure 3:

Difference (relative to WT) in IPOD-HR occupancies and difference (relative to WT) in RNA polymerase ChIP-seq rz-scores in indicated genotypes at individual replicate levels. For IPOD-HR, 50 bp rolling median was used as the fundamental unit of data for each genomic location, and the plotted values reflect the pseudomedian of those values across the indicated genomic features. Wilcoxon test was performed to obtain the error bars of the 95% confidence intervals. For RNA polymerase ChIP-seq, we followed a similar procedure as for IPOD-HR, except that we used gene-level means of the ChIP occupancies as individual units of data for the pseudomedian calculations.

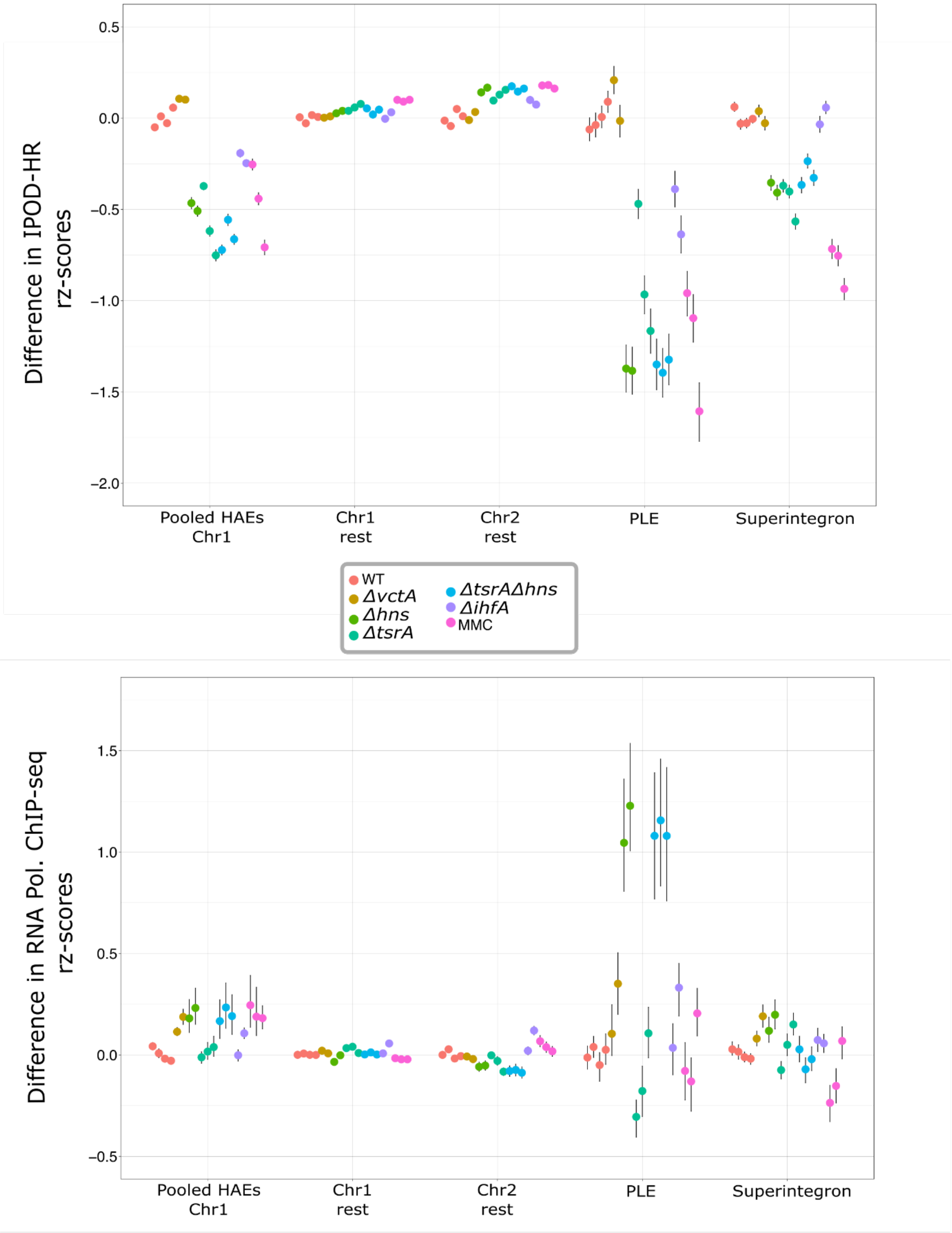

Supplementary Figure 4:

Heatmaps representing the differences of means of A) IPOD, B) IPOD-HR and C) RNA pol ChIP-seq rz-scores in the indicated conditions at biological replicate level from wild type.

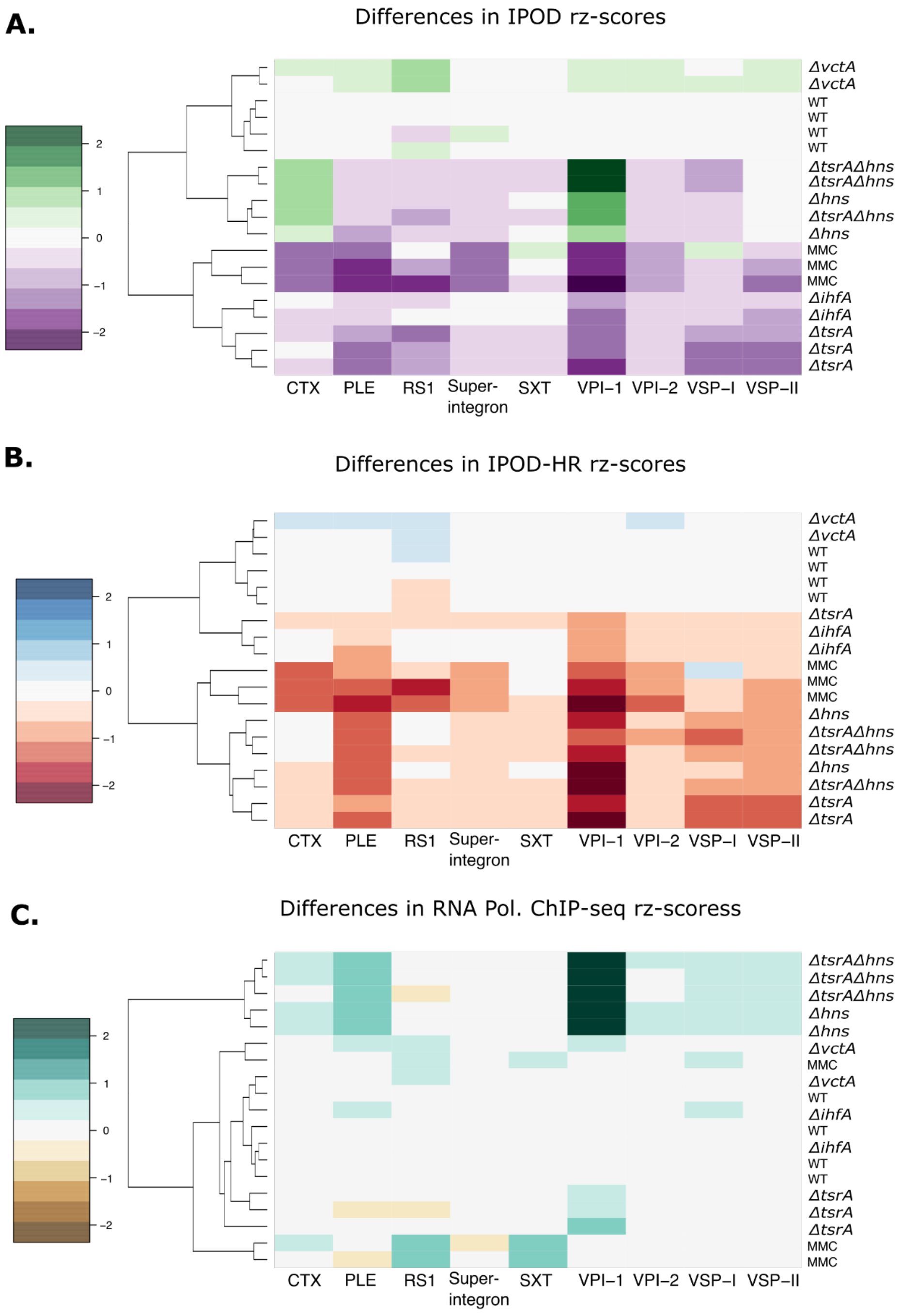

Supplementary figure 5 (next page):

A) Representing IPOD-HR tracks of the VPI-1 island for the indicated strains.
B) Distribution of scores in the wild type and ∆*ihfA* Vibrio cholerae KDS1 in the regions obtained from V5-H-NS ChIP-seq study in C6706 strain [71]
C) Representation of IPOD, IPOD-HR and RNA polymerase ChIP-seq tracks for all of the genotypes in the CTX region of *V. cholerae,* with the gray track being the V5-H-NS ChIP-seq from the C6706 study [71].
D) Representation of IPOD, IPOD-HR and RNA polymerase ChIP-seq tracks for all of the genotypes in the PLE region of *V. cholerae.*

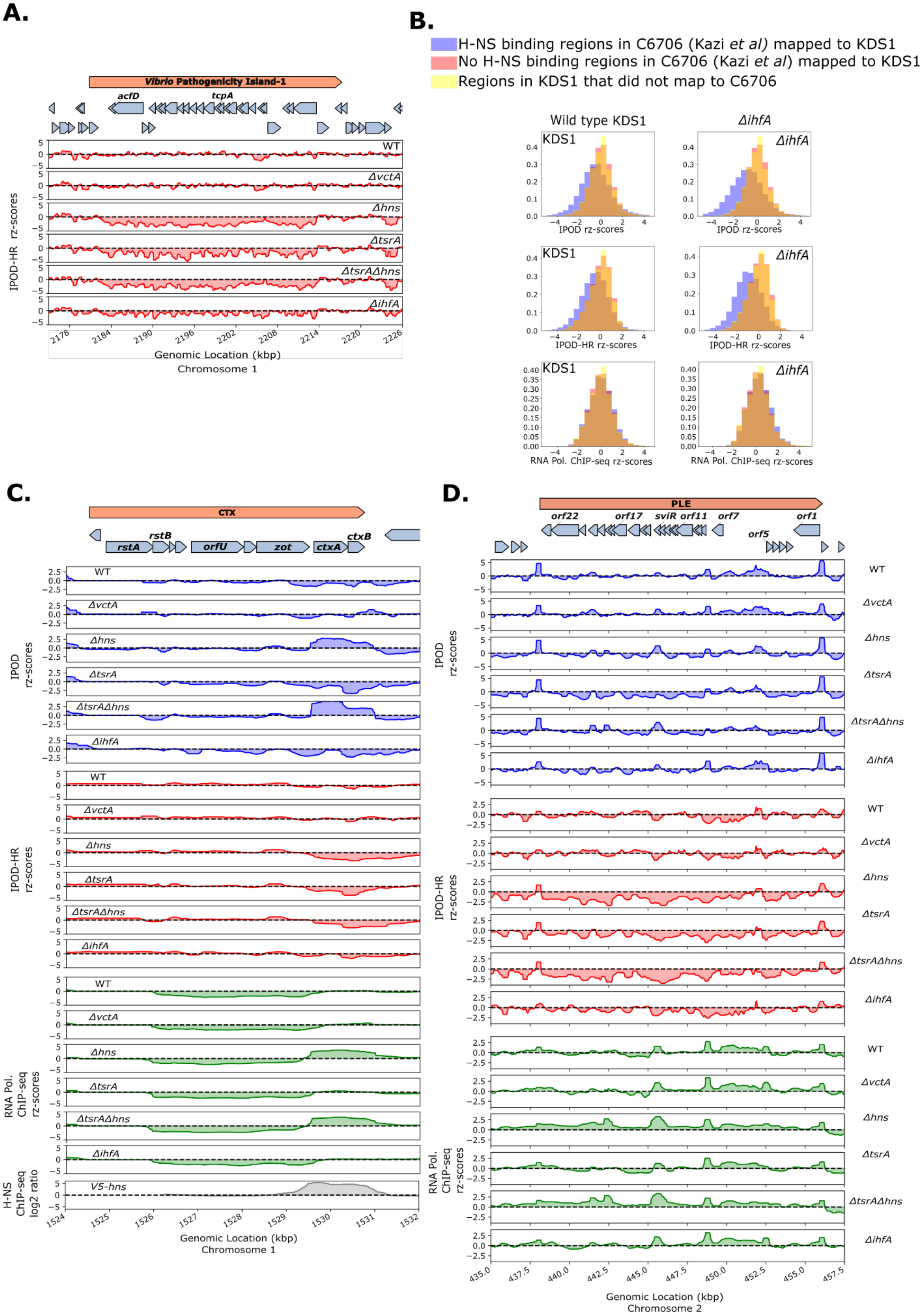
A.
Vibrio Pathogenicity Island-1
acfD
tcpA
WT
ΔvctA
Δhns
ΔtsrA
ΔtsrAΔhns
ΔihfA
IPOD-HR rz-scores
Genomic Location (kbp)
Chromosome 1
B.
H-NS binding regions in C6706 (Kazi et al) mapped to KDS1
No H-NS binding regions in C6706 (Kazi et al) mapped to KDS1
Regions in KDS1 that did not map to C6706
Wild type KDS1
ΔihfA
KDS1
IPOD rz-scores
IPOD-HR rz-scores
RNA Pol. ChIP-seq rz-scores
C.
CTX
rstB
ctxB
rstA
orfU
zot
ctxA
IPOD rz-scores
IPOD-HR rz-scores
RNA Pol. ChIP-seq rz-scores
H-NS ChIP-seq log2 ratio
V5-hns
Genomic Location (kbp)
Chromosome 1
D.
PLE
orf22
orf17
sviR
orf11
orf7
orf5
orf1
IPOD rz-scores
IPOD-HR rz-scores
RNA Pol. ChIP-seq rz-scores
Genomic Location (kbp)
Chromosome 2

Supplementary figure 6 (next page):

A) GO term enrichment classification analysis of RNA-sequencing results of indicated genotypes: ∆*hns,* ∆*tsrA,* and ∆*hns*∆*tsrA.*

B) AT percentage of differentially expressed genes in ∆*hns,* ∆*tsrA,* and ∆*hns*∆*tsrA* versus wild type. Significant differentially expressed genes (q-value less than or equal to 0.1 are indicated in black and q-value of greater than 0.1 are indicated in gray).

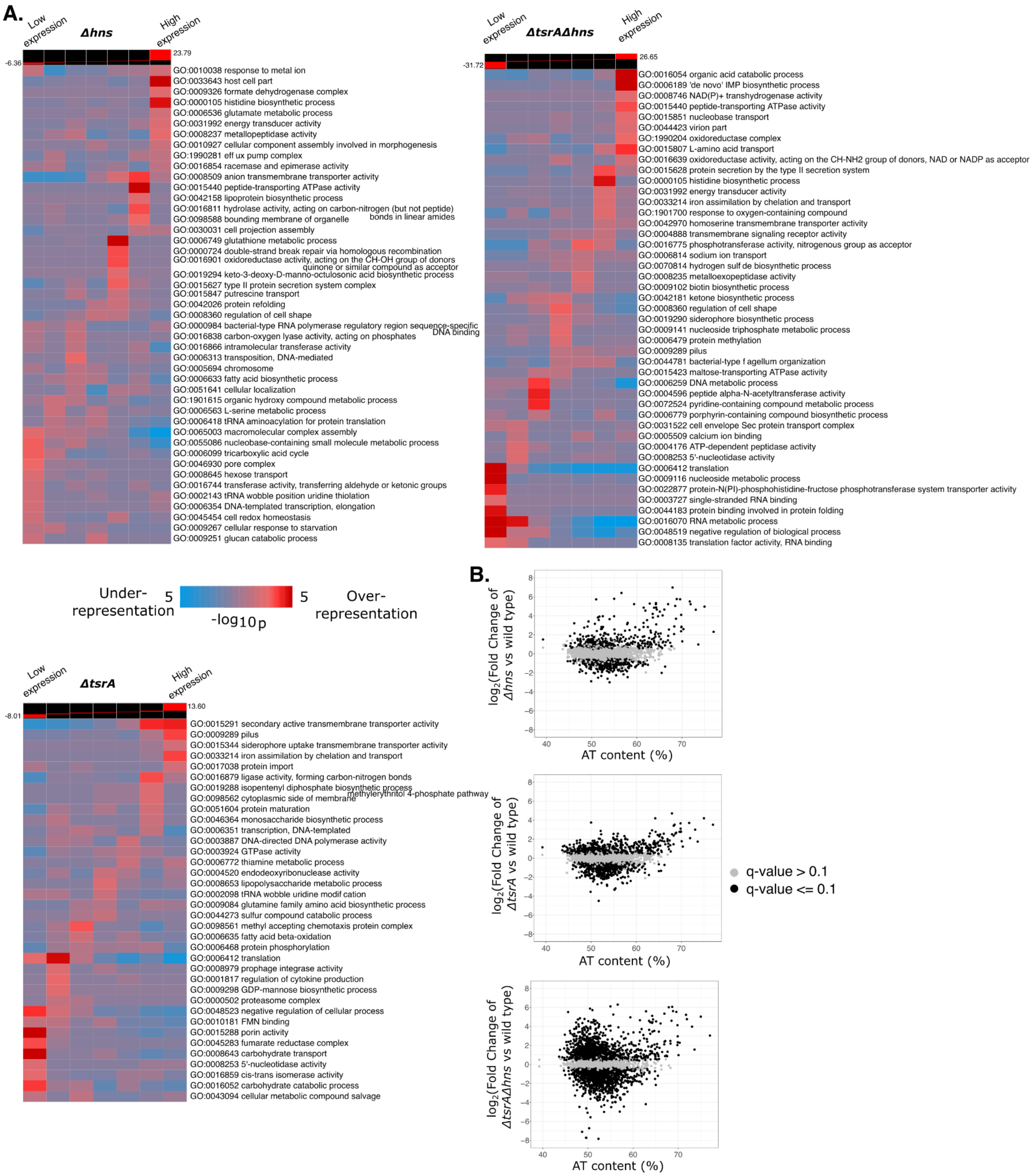

A.
Low expression
Δhns
High expression
-6.36
23.79
GO:0010038 response to metal ion
GO:0033643 host cell part
GO:0009326 formate dehydrogenase complex
GO:0000105 histidine biosynthetic process
GO:0006536 glutamate metabolic process
GO:0031992 energy transducer activity
GO:0008237 metallopeptidase activity
GO:0010927 cellular component assembly involved in morphogenesis
GO:1990281 eff ux pump complex
GO:0016854 racemase and epimerase activity
GO:0008509 anion transmembrane transporter activity
GO:0015440 peptide-transporting ATPase activity
GO:0042158 lipoprotein biosynthetic process
GO:0016811 hydrolase activity, acting on carbon-nitrogen (but not peptide) bonds in linear amides
GO:0098588 bounding membrane of organelle
GO:0030031 cell projection assembly
GO:0006749 glutathione metabolic process
GO:0000724 double-strand break repair via homologous recombination
GO:0016901 oxidoreductase activity, acting on the CH-OH group of donors quinone or similar compound as acceptor
GO:0019294 keto-3-deoxy-D-manno-octulosonic acid biosynthetic process
GO:0015627 type II protein secretion system complex
GO:0015847 putrescine transport
GO:0042026 protein refolding
GO:0008360 regulation of cell shape
GO:0000984 bacterial-type RNA polymerase regulatory region sequence-specific DNA binding
GO:0016838 carbon-oxygen lyase activity, acting on phosphates
GO:0016866 intramolecular transferase activity
GO:0006313 transposition, DNA-mediated
GO:0005694 chromosome
GO:0006633 fatty acid biosynthetic process
GO:0051641 cellular localization
GO:1901615 organic hydroxy compound metabolic process
GO:0006563 L-serine metabolic process
GO:0006418 tRNA aminoacylation for protein translation
GO:0065003 macromolecular complex assembly
GO:0055086 nucleobase-containing small molecule metabolic process
GO:0006099 tricarboxylic acid cycle
GO:0046930 pore complex
GO:0008645 hexose transport
GO:0016744 transferase activity, transferring aldehyde or ketonic groups
GO:0002143 tRNA wobble position uridine thiolation
GO:0006354 DNA-templated transcription, elongation
GO:0045454 cell redox homeostasis
GO:0009267 cellular response to starvation
GO:0009251 glucan catabolic process
Low expression
ΔtsrAΔhns
High expression
-31.72
26.65
GO:0016054 organic acid catabolic process
GO:0006189 'de novo' IMP biosynthetic process
GO:0008746 NAD(P)+ transhydrogenase activity
GO:0015440 peptide-transporting ATPase activity
GO:0015851 nucleobase transport
GO:0044423 virion part
GO:1990204 oxidoreductase complex
GO:0015807 L-amino acid transport
GO:0016639 oxidoreductase activity, acting on the CH-NH2 group of donors, NAD or NADP as acceptor
GO:0015628 protein secretion by the type II secretion system
GO:0000105 histidine biosynthetic process
GO:0031992 energy transducer activity
GO:0033214 iron assimilation by chelation and transport
GO:1901700 response to oxygen-containing compound
GO:0042970 homoserine transmembrane transporter activity
GO:0004888 transmembrane signaling receptor activity
GO:0016775 phosphotransferase activity, nitrogenous group as acceptor
GO:0006814 sodium ion transport
GO:0070814 hydrogen sulf de biosynthetic process
GO:0008235 metalloexopeptidase activity
GO:0009102 biotin biosynthetic process
GO:0042181 ketone biosynthetic process
GO:0008360 regulation of cell shape
GO:0019290 siderophore biosynthetic process
GO:0009141 nucleoside triphosphate metabolic process
GO:0006479 protein methylation
GO:0009289 pilus
GO:0044781 bacterial-type f agellum organization
GO:0015423 maltose-transporting ATPase activity
GO:0006259 DNA metabolic process
GO:0004596 peptide alpha-N-acetyltransferase activity
GO:0072524 pyridine-containing compound metabolic process
GO:0006779 porphyrin-containing compound biosynthetic process
GO:0031522 cell envelope Sec protein transport complex
GO:0005509 calcium ion binding
GO:0004176 ATP-dependent peptidase activity
GO:0008253 5'-nucleotidase activity
GO:0006412 translation
GO:0009116 nucleoside metabolic process
GO:0022877 protein-N(PI)-phosphohistidine-fructose phosphotransferase system transporter activity
GO:0003727 single-stranded RNA binding
GO:0044183 protein binding involved in protein folding
GO:0016070 RNA metabolic process
GO:0048519 negative regulation of biological process
GO:0008135 translation factor activity, RNA binding
Under-representation
5
5
Over-representation
-log10p
Low expression
ΔtsrA
High expression
-8.01
13.60
GO:0015291 secondary active transmembrane transporter activity
GO:0009289 pilus
GO:0015344 siderophore uptake transmembrane transporter activity
GO:0033214 iron assimilation by chelation and transport
GO:0017038 protein import
GO:0016879 ligase activity, forming carbon-nitrogen bonds
GO:0019288 isopentenyl diphosphate biosynthetic process methylerythritol 4-phosphate pathway
GO:0098562 cytoplasmic side of membrane
GO:0051604 protein maturation
GO:0046364 monosaccharide biosynthetic process
GO:0006351 transcription, DNA-templated
GO:0003887 DNA-directed DNA polymerase activity
GO:0003924 GTPase activity
GO:0006772 thiamine metabolic process
GO:0004520 endodeoxyribonuclease activity
GO:0008653 lipopolysaccharide metabolic process
GO:0002098 tRNA wobble uridine modif cation
GO:0009084 glutamine family amino acid biosynthetic process
GO:0044273 sulfur compound catabolic process
GO:0098561 methyl accepting chemotaxis protein complex
GO:0006635 fatty acid beta-oxidation
GO:0006468 protein phosphorylation
GO:0006412 translation
GO:0008979 prophage integrase activity
GO:0001817 regulation of cytokine production
GO:0009298 GDP-mannose biosynthetic process
GO:0000502 proteasome complex
GO:0048523 negative regulation of cellular process
GO:0010181 FMN binding
GO:0015288 porin activity
GO:0045283 fumarate reductase complex
GO:0008643 carbohydrate transport
GO:0008253 5'-nucleotidase activity
GO:0016859 cis-trans isomerase activity
GO:0016052 carbohydrate catabolic process
GO:0043094 cellular metabolic compound salvage
B.
log2(Fold Change of Δhns vs wild type)
AT content (%)
log2(Fold Change of ΔtsrA vs wild type)
AT content (%)
q-value > 0.1
q-value <= 0.1
log2(Fold Change of ΔtsrAΔhns vs wild type)
AT content (%)

Supplementary Figure 7 (next page):

GO term enrichment classification analysis of the upregulated (three times the standard error of the significantly differentially expressed genes (q-value less than or equal to 0.1) compared to wild type) genes from the RNA-sequencing results of indicated genotypes: ∆*hns,* ∆*tsrA,* and ∆*hns*∆*tsrA.*

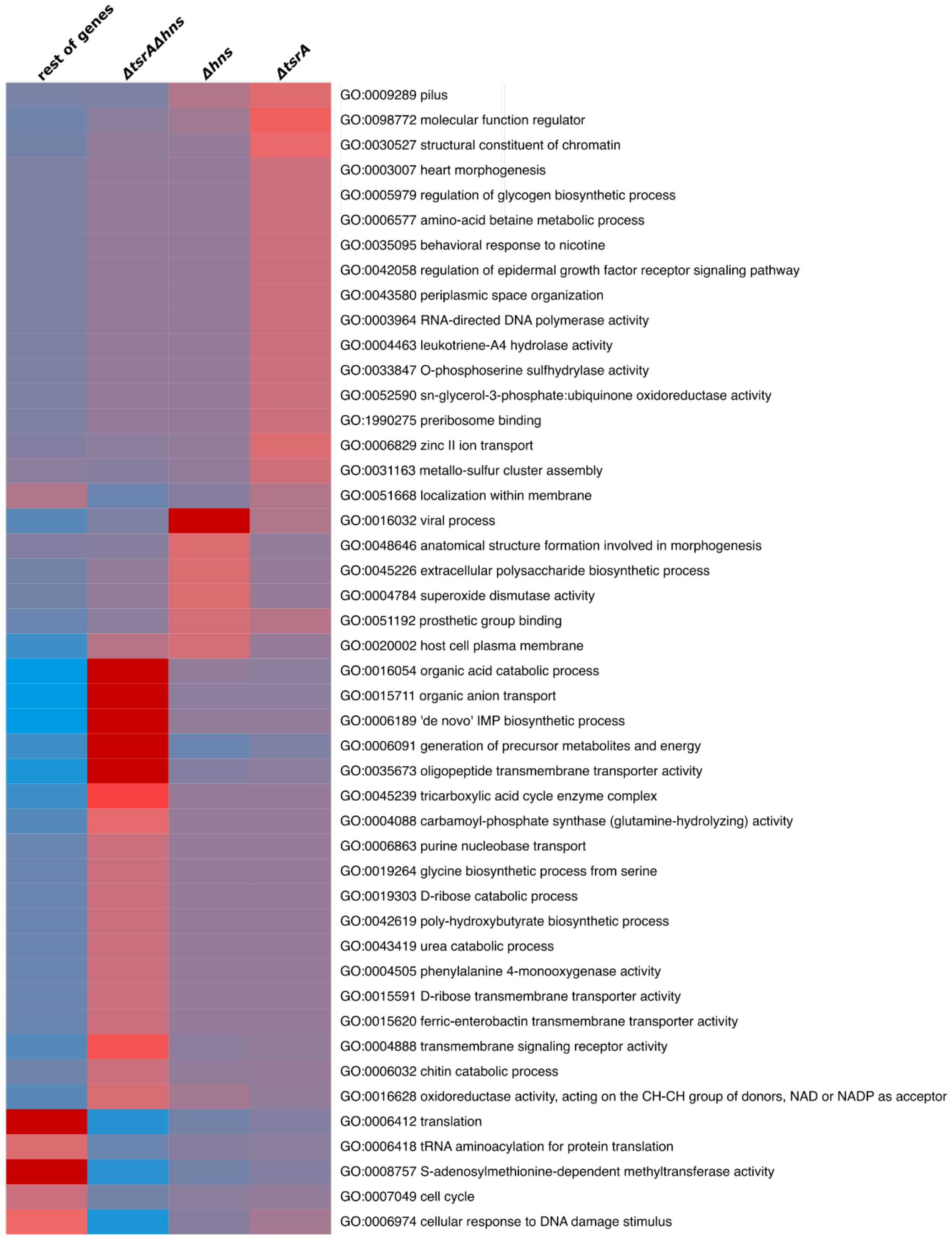
rest of genes
ΔtsrAΔhns
Δhns
ΔtsrA
GO:0009289 pilus
GO:0098772 molecular function regulator
GO:0030527 structural constituent of chromatin
GO:0003007 heart morphogenesis
GO:0005979 regulation of glycogen biosynthetic process
GO:0006577 amino-acid betaine metabolic process
GO:0035095 behavioral response to nicotine
GO:0042058 regulation of epidermal growth factor receptor signaling pathway
GO:0043580 periplasmic space organization
GO:0003964 RNA-directed DNA polymerase activity
GO:0004463 leukotriene-A4 hydrolase activity
GO:0033847 O-phosphoserine sulfhydrylase activity
GO:0052590 sn-glycerol-3-phosphate:ubiquinone oxidoreductase activity
GO:1990275 preribosome binding
GO:0006829 zinc II ion transport
GO:0031163 metallo-sulfur cluster assembly
GO:0051668 localization within membrane
GO:0016032 viral process
GO:0048646 anatomical structure formation involved in morphogenesis
GO:0045226 extracellular polysaccharide biosynthetic process
GO:0004784 superoxide dismutase activity
GO:0051192 prosthetic group binding
GO:0020002 host cell plasma membrane
GO:0016054 organic acid catabolic process
GO:0015711 organic anion transport
GO:0006189 'de novo' IMP biosynthetic process
GO:0006091 generation of precursor metabolites and energy
GO:0035673 oligopeptide transmembrane transporter activity
GO:0045239 tricarboxylic acid cycle enzyme complex
GO:0004088 carbamoyl-phosphate synthase (glutamine-hydrolyzing) activity
GO:0006863 purine nucleobase transport
GO:0019264 glycine biosynthetic process from serine
GO:0019303 D-ribose catabolic process
GO:0042619 poly-hydroxybutyrate biosynthetic process
GO:0043419 urea catabolic process
GO:0004505 phenylalanine 4-monooxygenase activity
GO:0015591 D-ribose transmembrane transporter activity
GO:0015620 ferric-enterobactin transmembrane transporter activity
GO:0004888 transmembrane signaling receptor activity
GO:0006032 chitin catabolic process
GO:0016628 oxidoreductase activity, acting on the CH-CH group of donors, NAD or NADP as acceptor
GO:0006412 translation
GO:0006418 tRNA aminoacylation for protein translation
GO:0008757 S-adenosylmethionine-dependent methyltransferase activity
GO:0007049 cell cycle
GO:0006974 cellular response to DNA damage stimulus

Supplementary Figure 8:

A) Correlation plot of rz-log ratios of RNA polymerase ChIP-seq vs. Input with log10(Transcripts per million (TPM)) in the wild type KDS1. The data are of the mean of four replicates from each experiment: RNA pol. ChIP-seq and RNA-seq. The line represents a robust linear model of Log10(TPM) as a function of RNA polymerase ChIP-seq in wild type. The slope is shown in blue. Genes with less than 0.0001 TPM are omitted.
B) Occupancy traces of IPOD, IPOD-HR and RNA polymerase ChIP-seq of the whole SXT-VchInd6 genomic feature in untreated and MMC treated wild type *Vibrio cholerae*. The shaded rectangles are shown for visual comparison of regions with differences in RNA polymerase binding between the untreated and MMC treated cells.

**A.**

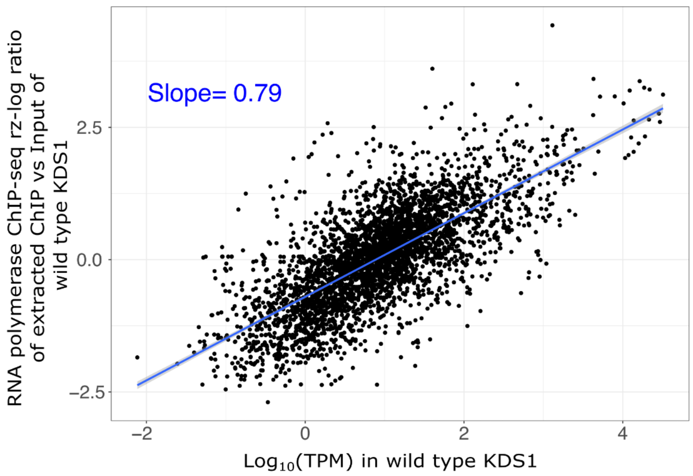


**B.**

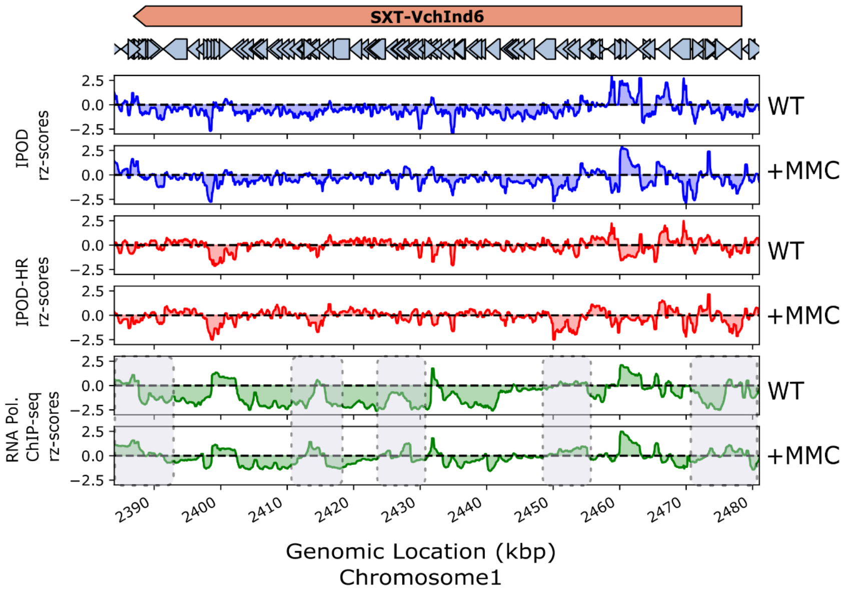

Supplementary Table 1: Numbers of overlapping and unique peaks in the conditions tested compared to the 122 peaks in the wild type.

| Strain | Number of peaks | Overlapping peaks | Unique peaks |
|---|---|---|---|
| Δ*ihf* | 106 | 38 | 68 |
| Δ*hns* | 100 | 52 | 48 |
| Δ*tsrA* | 98 | 54 | 44 |
| Δ*vctA* | 116 | 64 | 52 |
| Δ*tsrA*Δ*hns* | 121 | 42 | 79 |
| MMC | 33 | 24 | 9 |

Supplementary Table 2: Bacterial strains

| Strain and genotype | Source | Lab identifier |
|---|---|---|
| *V. cholerae* PLE(+) | Seed et al., 2013 [55] | KDS1 |
| *V. cholerae* PLE(+)Δ*hns*::frt-spec-frt | This study | DD427 |
| *V. cholerae* PLE(+)Δ*ihfA* | This study | DD430 |
| *V. cholerae* PLE(+)Δ*tsrA*::frt-kan-frt | This study | DD601 |
| *V. cholerae* PLE(+)Δ*tsrA*::frt-kan-frt + Δ*hns*::frt-spec-frt | This study | DD607 |
| *V. cholerae* PLE(+)Δ*vctA* | This study | KS2599 |
